# Generative and predictive neural networks for the design of functional RNA molecules

**DOI:** 10.1101/2023.07.14.549043

**Authors:** Aidan T. Riley, James M. Robson, Alexander A. Green

**Affiliations:** Department of Biomedical Engineering, Boston University, Boston, MA 02215, USA; Biological Design Center, Boston University, Boston, MA 02215, USA; Molecular Biology, Cell Biology & Biochemistry Program, Graduate School of Arts and Sciences, Boston University, Boston, MA 02215, USA

## Abstract

RNA is a remarkably versatile molecule that has been engineered for applications in therapeutics, diagnostics, and *in vivo* information-processing systems. However, the complex relationship between the sequence and structural properties of an RNA molecule and its ability to perform specific functions often necessitates extensive experimental screening of candidate sequences. Here we present a generalized neural network architecture that utilizes the <u>s</u>equence <u>and s</u>tructure <u>o</u>f <u>R</u>NA <u>m</u>olecules (SANDSTORM) to inform functional predictions. We demonstrate that this approach achieves state-of-the-art performance across several distinct RNA prediction tasks, while learning interpretable abstractions of RNA secondary structure. We paired these predictive models with <u>g</u>enerative <u>a</u>dversarial <u>R</u>NA <u>d</u>esign <u>n</u>etworks (GARDN), allowing the generative modelling of novel mRNA 5’ untranslated regions and toehold switch riboregulators exhibiting a predetermined fitness. This approach enabled the design of novel toehold switches with a 43-fold increase in experimentally characterized dynamic range compared to those designed using classic thermodynamic algorithms. SANDSTORM and GARDN thus represent powerful new predictive and generative tools for the development of diagnostic and therapeutic RNA molecules with improved function.

## INTRODUCTION

The ability to encode genetic information while executing a diverse range of functions have allowed RNA to gain traction as a powerful molecular platform for the development of human therapeutics, diagnostics, and synthetic biological systems^1–11^. The quartet of canonical nucleotides and simple rules of Watson-Crick base pairing make the design of specific RNA molecules a far more computationally tractable problem than analogous protein-based systems. Extensive work has been conducted to develop robust tools for the prediction of RNA secondary structure and structure-based inverse design^12–18^. However, there does not exist an equally generalized set of tools for computational engineering of RNA function. As RNA-based systems have become more sophisticated and widely used, a disconnect between deterministic thermodynamic properties and resulting RNA function has grown, often necessitating experimental screening of many RNA sequences^3, 5, 7, 9, 11^,. Accordingly, data-driven methods that can decipher the relationship between RNA sequence and function are essential for expediting RNA device design and improving their performance in biomedical applications.

To facilitate data-based approaches to RNA engineering, massively parallel reporting assays have been designed across a variety of contexts^19–32^, resulting in a number of datasets mapping functional RNAs to their experimental performance. Toehold switches^33, 34^, 5’ untranslated regions (UTRs)^35^, CRISPR guide RNAs^36^, and ribosome binding sites (RBSs)^37^ are all examples of RNA sequences that have been functionally characterized using high-throughput screening. In each of these cases, predictive models that operate on a one-hot-encoded representation of the input sequences have been constructed alongside the datasets. The architectures and performance of these models vary widely, despite their common task of predicting RNA function. This has resulted in several available predictive methods that are tuned extensively to their specific functional task but cannot be applied to the full span of settings in which RNA has found use. Just as thermodynamic algorithms for sequence-structure relationships can be applied regardless of the class of RNA being analyzed, we aimed to develop a similarly generalized framework for predicting RNA sequence-function relationships and designing new instances of RNA molecules with a pre-determined function. A robust set of tools that enable the prediction of RNA function and the inverse design of functional sequences could greatly accelerate the deployment of RNA systems in both research and industrial settings.

Despite both sequence and structure playing a large role in the function of RNA molecules, there currently does not exist a generalized deep learning framework that incorporates both properties to inform predictions. Here we report a deep learning approach that incorporates the <u>s</u>equence <u>and st</u>ructure <u>o</u>f <u>R</u>NA <u>m</u>olecules (SANDSTORM) to predict the function of diverse classes of RNAs. Using SANDSTORM neural networks, we predict the functional activity of 5’ UTRs, ribosome binding sites, CRISPR guide RNAs, and toehold switch riboregulators. By combining these predictive models with <u>G</u>enerative <u>A</u>dversarial <u>R</u>NA <u>D</u>esign <u>N</u>etworks (GARDN), we generate novel and realistic instances of several of these RNA classes that also exhibit desirable functional properties. This joint design method demonstrates that the abstractions of secondary structure learned by the SANDSTORM architecture can be effectively leveraged by the generator during training, resulting in generated constructs that reflect experimentally characterized thermodynamic properties. The ability to both predict and generate RNA sequences with tailored functional activity could serve as the foundation of a wide range of next-generation computational RNA tools.

## RESULTS

### Implementing an efficient representation of RNA secondary structure

In order to predict function using both sequence and structure, we first developed a representation of RNA secondary structure that could be supplied to a predictive model in parallel with one-hot-encoded sequences (Fig. 1). To do so, we took inspiration from the field of image classification, where manually engineered data features have been surpassed by deep convolutional models which learn their own abstractions of input data^38^. By constructing a basic array that encoded only the locations of possible base pairs and the distance between these bases, we hypothesized that a deep CNN would be able to construct similar abstractions related to the structure of an RNA molecule during the training process. The novel structural array we present is thematically similar to others that have been developed for RNA inverse design^16, 33^, but offers both the time and memory efficiency necessary to be practically useful in a deep learning pipeline while avoiding any assumptions inherent to classic structural prediction algorithms (Supplementary Fig. 2).

**Figure 1.**
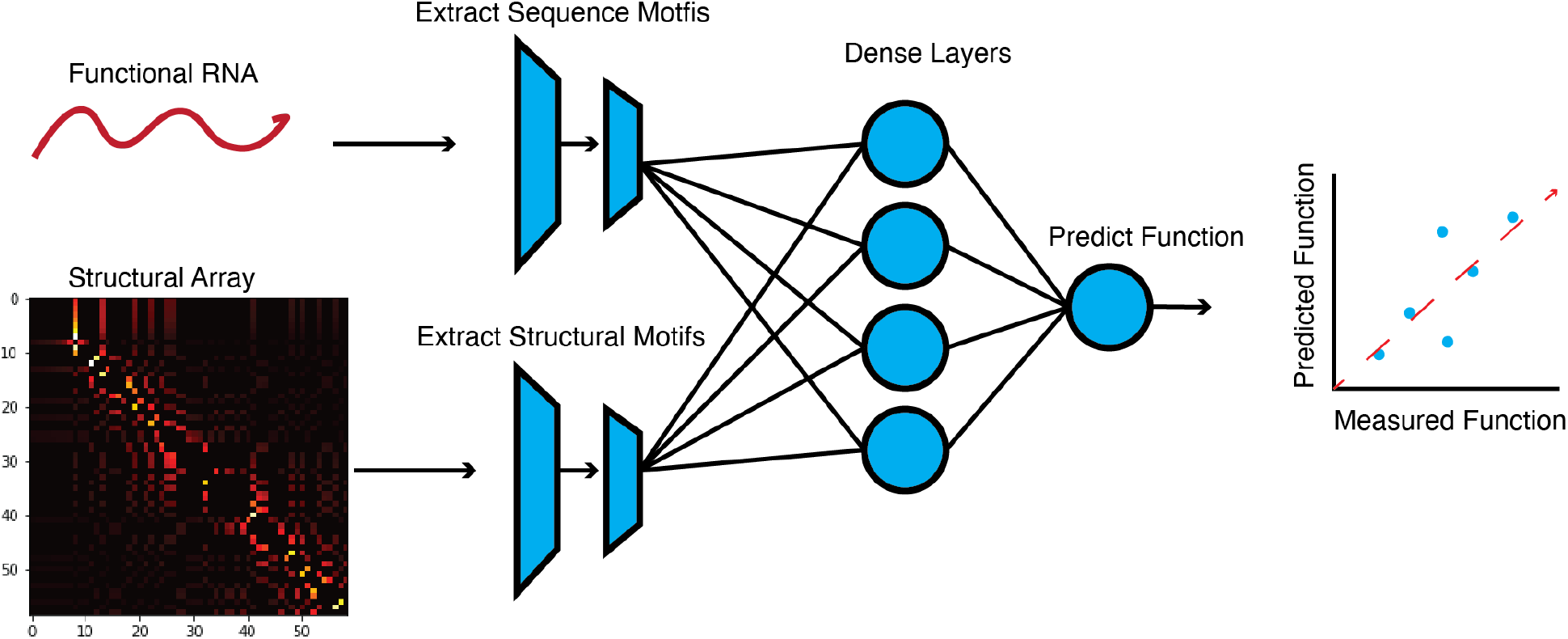
Overview of the SANDSTORM RNA prediction model. SANDSTORM expands upon previous sequence-to-function neural networks by incorporating both sequence and secondary structure array input channels. These paired inputs are passed through parallel convolutional stacks that form an ensemble prediction of input RNA function (see Supplementary Fig. 1 for a detailed depiction of SANDSTORM).

### Learning interpretable structural information with a simulated dataset

We evaluated a CNN’s ability to convert our structural array into interpretable structural information using a simulated dataset of toehold switches and several decoy sequences (Fig. 2a). Toehold switches are riboregulators that modulate translation of an output protein in response to a complementary RNA input molecule^3^. These devices are designed to hide the RBS and start codon from the translational machinery by placing them within a cis-repressing stem. This stem can be unwound by an RNA trigger that is complementary to the 5’ end of the switch, releasing both the RBS and start codon. Because these devices require both sequence and structural motifs to function, they serve as an ideal candidate to demonstrate the performance improvement realized by using both properties as inputs. A dataset was constructed consisting of canonical toehold switches and four types of decoy sequences: sequences containing only an RBS motif and random nucleotides; sequences containing only a start codon motif; sequences containing both a start codon and RBS, but do not fold into the toehold switch structure; and sequences that fold into the toehold switch structure, but do not contain the necessary sequence-level motifs.

**Figure 2.**
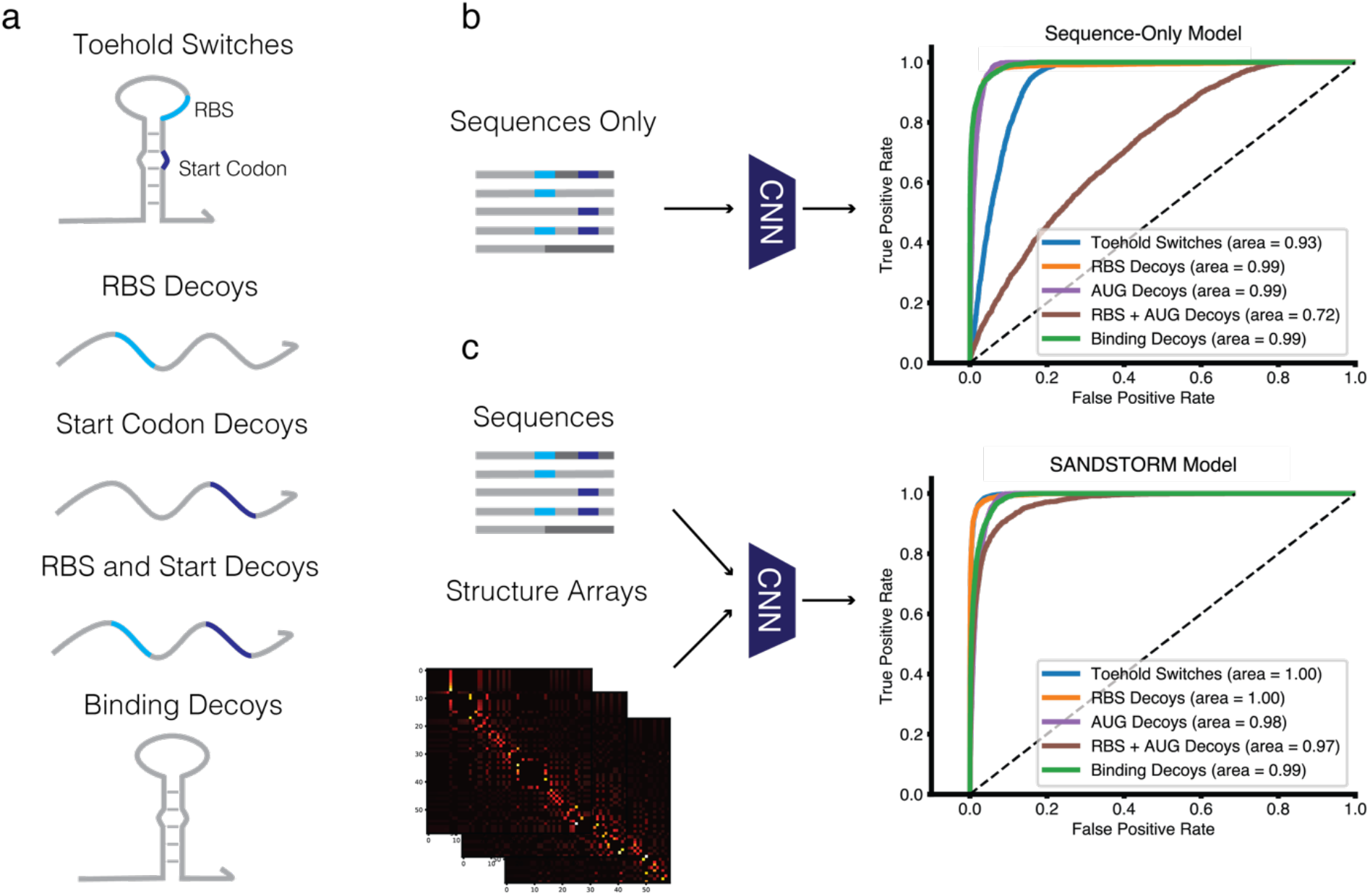
Extracting secondary structure information using a simulated dataset. **a**, A dataset of toehold switches and several types of decoy sequences was utilized to determine whether a CNN could differentiate sequences on a structural basis. Sequences consisted of canonical toehold switches, decoys containing only an RBS motif (RBS decoys), decoys containing only a start codon motif (AUG decoys), decoys containing both an RBS and start codon motif (RBS + AUG decoys), and decoys that adopted the canonical secondary structure but did not contain the necessary sequence motifs (binding decoys). **b**, A CNN trained only on one-hot-encoded sequences was not able to perfectly classify the canonical toehold switches from the RBS + AUG decoys, which are only differentiable at a structural level. **c**, The SANDSTORM CNN accepting paired sequence and structure arrays was able to identify both the sequence and structural features required to classify the simulated dataset.

We compared the performance of a model accepting only one-hot-encoded sequences as inputs to the SANDSTORM model accepting a paired sequence-structure input at the task of classifying each member of this simulated dataset (Fig. 2b,c). As expected, the one-hot-encoding model did not perform as well as the dual-input model when differentiating between canonical toehold switches and decoys that contained the necessary sequence motifs but did not fold into the correct secondary structure (single-input sequence-only model AUC=0.72, SANDSTORM model AUC=0.97). As these two categories are only differentiable at the secondary structure level, we concluded that our novel array enabled the model to learn useful abstractions of RNA structure to inform predictions.

### SANDSTORM models predict the function of multiple classes of RNAs

After demonstrating the capability of the SANDSTORM architecture to account for secondary structure, we sought to determine if this approach could be generalized to predict the function of different classes of real RNA molecules (Fig. 3). As our goal was to develop a generalized framework for predicting RNA function, we designed a single SANDSTORM CNN architecture that could be applied to any RNA prediction task without sacrificing performance. Our model consisted of a sequence input channel and a structure input channel, each connected to an independent stack of convolutional layers. The outputs of the convolutional channels were concatenated before being passed through a final series of densely connected layers to allow the model to learn complex relationships between the features abstracted from each input. To avoid overfitting to a specific task, the model consisted of as few trainable parameters as possible and the hyperparameters were not tuned for each presented test case. The same data split was used to compare the mean squared error (MSE), R^2^, and Spearman correlation for SANDSTORM and the previously published model for each regression task.

**Figure 3.**
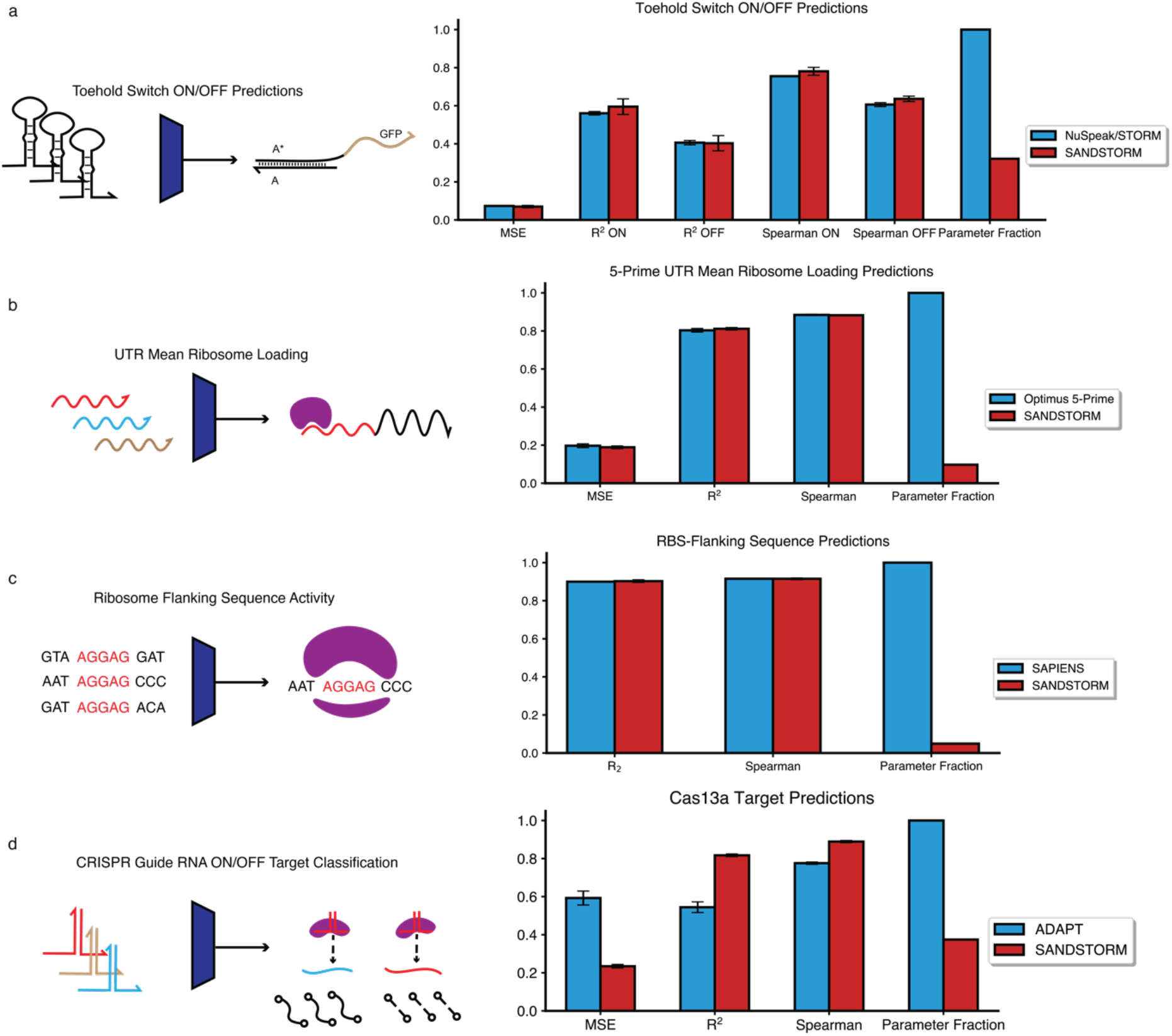
SANDSTORM models can predict the function of several classes of RNA molecules. **a**, SANDSTORM prediction metrics for toehold switch ON and OFF values compared to NuSpeak/STORM. **b**, SANDSTORM predictions for UTR mean ribosome loading compared to Optimus 5-prime. **c**, SANDSTORM model predicting RBS translation efficiency compared to SAPIENS. **d**, SANDSTORM predictions of Cas13a collateral cleavage efficiency using guide RNA-target pairs compared to ADAPT. Bars represent three-fold cross validation mean ± s.d.

#### Toehold switch prediction

Angenent-Mari et al.^33^ conducted an assay that sorted a large library of toehold switches based on their resulting fluorescence levels measured in both the native folding conformation, which corresponds to the OFF state, and when fused to the cognate trigger, which is equivalent to the ON state. This dataset therefore consists of toehold switch sequences that are mapped to a fluorescence readout in the ON and OFF states, with desirable sequences having the highest ON/OFF ratio possible. Despite the wealth of data mapping sequence to function for toehold switches, there is currently no theoretical framework that can reliably link sequence or thermodynamic characteristics to ON/OFF ratios. Furthermore, previous studies attempting to use engineered thermodynamic features as the inputs to a multi-layer-perceptron similarly failed to produce strong predictions^33^. The SANDSTORM-toehold architecture can match the most successful model from previous studies^33–34^ (cross-fold MSE p-value = 0.43, two-sided t-test) values while using 67% fewer trainable parameters (Fig. 3a).

#### 5’ UTR prediction

Several cis-regulatory elements affect the translational efficiency of full-length messenger RNAs in eukaryotic cells. One such element that can have a profound effect on protein production is the 5’ UTR; however, a detailed prediction of the regulatory effect cannot be captured by first-order sequence or structural relationships^39–40^. To survey the space of regulatory UTRs, Sample et al.^35^ conducted a high-throughput polysome profiling assay and constructed a predictive CNN using the resulting sequence-function dataset. Our SANDSTORM-UTR architecture was able to match the previously reported performance metrics (cross fold MSE p-value = 0.059, two-sided t-test) on this dataset (Fig. 3b) while using only 9.7% of the number of trainable parameters, suggesting that the predictive capabilities of SANDSTORM models can be applied across RNA classes.

#### RBS-flanking sequence prediction

The RBS sequence motif is required for translation initiation in prokaryotic systems. While the core motif is necessary for translation to occur, the variable nucleotides surrounding the RBS can have a large impact on the resulting degree of protein production^11, 41^. This effect was quantified in a high-throughput manner by promoting translation of a site-specific DNA recombinase off a library of different RBS-flanking sequences^37^. When translated, this recombinase can flip the orientation of downstream section of the DNA molecule, allowing the fraction of DNA readouts that are flipped after sequencing to serve as a proxy for translation efficiency. As with the other cases reported here, the relationship between RBS-flanking sequences and translation efficiency was originally modelled using a sequence-only probabilistic deep learning framework. The SANDSTORM-RBS model was able to again match the predictive performance of the previously published approach (cross fold R^2^ p-value = 0.49, cross fold Spearman p-value=0.95, two-sided t-test) while using 95% fewer trainable parameters (Fig. 3c). As the original model published with this dataset used a probabilistic loss instead of mean squared error, the performance comparison consists of only the correlations between predicted and ground-truth outputs.

#### CRISPR Cas13a target-guide prediction

CRISPR/Cas nucleases enable site-specific cleavage of nucleic acids based on the spacer sequence supplied by a guide RNA^2^. Predicting the efficiency of on- and off-target sequences that could be cleaved by Cas nucleases is a major challenge in the development of CRISPR systems and has been approached with a variety of different methods. To tackle this challenge, high-throughput methods^42^ have been developed to map the relationship between gRNA/target RNA pairs and resulting Cas13a collateral cleavage efficiency. The SANDSTORM-Cas model is capable of significantly improving on the performance of the previously characterized sequence-only model^36^ (cross fold MSE p-value=5×10^−5^, two-sided t-test) while using only 37% of the trainable parameters (Fig. 3d). Additionally, this test case demonstrates that SANDSTORM architectures are capable of predicting RNA functions that are not tied directly to a translational readout, further supporting the generalizability of this approach.

### Implementing a generative architecture for RNA design

After establishing a generalized approach to predict RNA function, we aimed to develop a similarly robust framework for designing functional RNA sequences. While generative modelling has recently gained traction in the protein and DNA sequence design space^43–48^, there is a notable lack of these techniques applied to the RNA design task. Previous studies in the area of RNA design have deployed genetic algorithms to design single sequences which maximize the functional predictions of a pretrained CNN and activation maximization algorithms, where a single or paired set of sequence probability weight matrices are directly optimized based on the gradients of a predictive model^35, 47–50^. These approaches can notably suffer from low diversity in optimized sequences, as the same sequence motifs that maximize the predictive score will occur in the output regardless of the initial seed. Generative Adversarial Networks have demonstrated strikingly realistic and diverse results across several generative tasks^51–53^; however, their complexity can make it difficult to identify a stable training configuration. A design approach that can incorporate the realism offered by GANs with functional predictions has yet to be applied to the task of designing functional RNA molecules.

### GARDN for 5’ UTR Design

Here we generalize a GAN framework to construct novel nucleic acid sequences with a range of functions, beginning with 5’ UTRs regulating translation efficiency. Initially a generator and a discriminator model were trained under a standard Wasserstein GAN with gradient penalty (WGAN-GP) settings^52^ (Fig. 4a) until a generator was realized that could return examples of data that were indistinguishable from the real set. In the case of 5’ UTRs, the generator output is a single 50-nt long sequence represented by a 4×50 one-hot-encoded array. These arrays can be decoded back into the nucleotide letter space and their frequencies averaged to ensure that the generator is accurately modelling the nucleotide distribution of the real data (Fig. 4b).

**Figure 4.**
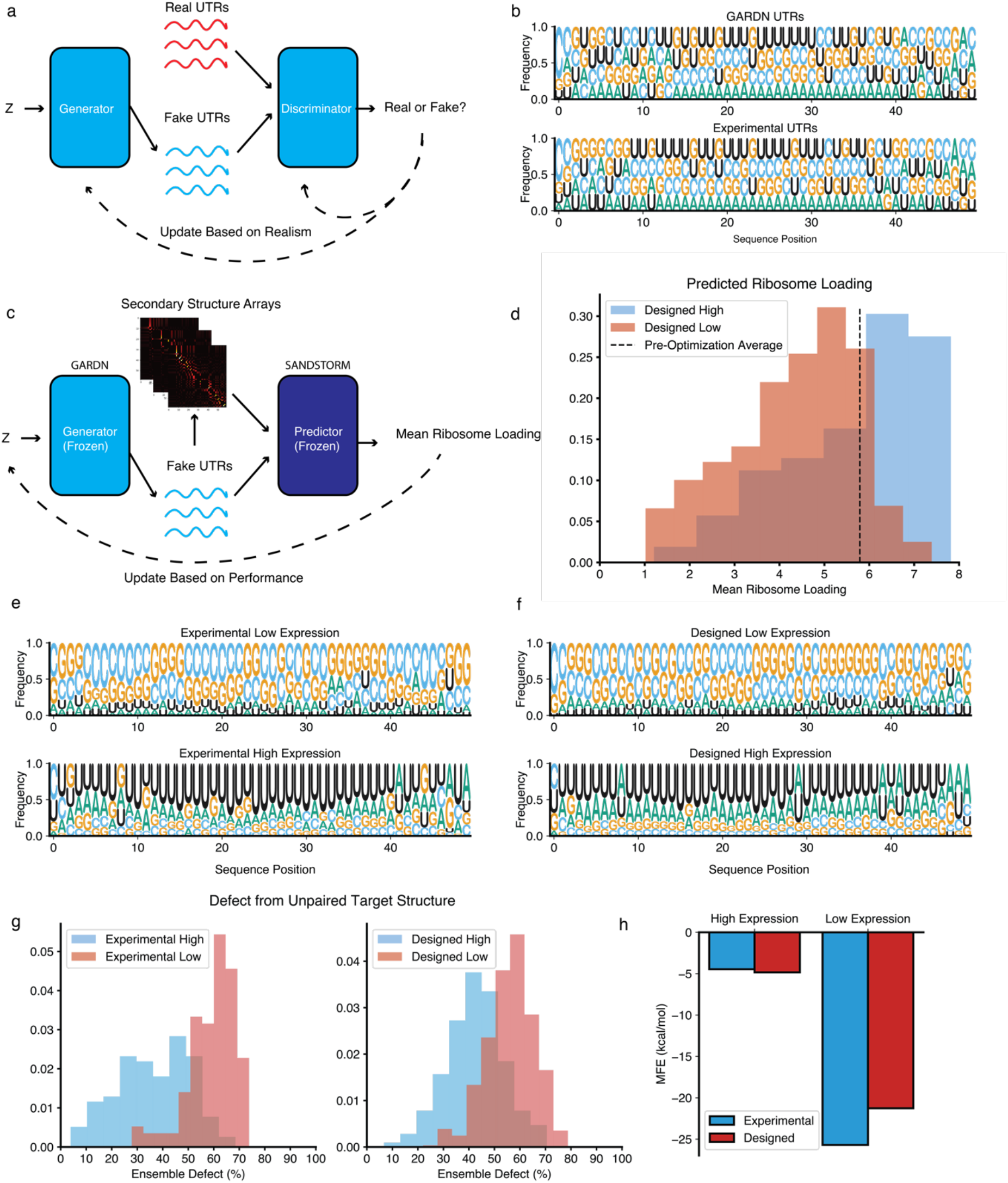
GARDN design of functional 5’ UTR sequences. **a**, An overview of the Generative Adversarial Network training setup, where a generator and predictor model are updated until realistic UTR sequences are returned. **b**, GARDN models can accurately model the nucleotide distribution of the real UTR dataset. **c**, Connecting a trained GARDN generator to a SANDSTORM-UTR predictor enables the design of realistic UTR sequences with desired functional properties. **d**, GARDN-generated UTRs can be pushed towards either predicted high or low expression. **e,f**, Sequence distributions of experimentally verified (e) and GARDN-designed (f) high- and low-expression UTRs. GARDN sequences match the distributions of the experimental dataset. **g**,**h** GARDN-generated UTRs show similar distributions of secondary structures compared to experimentally characterized UTRs, as measured by ensemble defect from a completely unpaired RNA structure (g) or minimum free energy (h).

Once we characterized a generator that could return realistic 5’ UTR samples, these sequences were fed into our SANDSTORM-UTR predictor model. By freezing the weights of both the generator and predictor, the latent space providing input seeds to the generator could be explored with respect to the downstream fitness score (Fig. 4c). An activation maximization algorithm linking latent inputs to fitness scores allowed us to identify regions in the latent space that generate sequences with desirable features. This approach is advantageous over direct activation maximization of a single sequence as the frozen generator enforces realism and diversity constraints on its outputs, even as the latent inputs are optimized to generate samples with extreme fitness scores. Furthermore, the lightweight architecture of the predictive models reduces the complexity of the optimization landscape, allowing improved sequences to be designed with minimal iterative calls to the predictor. Optimizing GARDN outputs enabled the generation of UTRs with both expected high and low translation efficiency (Fig. 4d).

When compared to the experimentally high- and low-performing UTRs, the GARDN-designed sequences optimized by the SANDSTORM-UTR model recapitulated not only the respective nucleotide distributions, but also thermodynamic properties known to affect translation efficiency for each category (Fig. 4e-h). Experimental evidence has shown that high translation efficiency is associated with minimal 5’ UTR secondary structure, while low translation efficiency is associated with extensive UTR secondary structure^37, 38, 48^. To interrogate whether our GARDN models recapitulated this trend, we calculated the ensemble defect of the UTRs from the completely unpaired target structure (Fig. 4g) as well as the minimum free energy (MFE) of the resulting sequences (Fig. 4h), demonstrating that the designed constructs recreated the thermodynamic distributions of their experimental targets. Additionally, we calculated the MFE of the designed UTRs in a sliding window over the length of each sequence to analyze if the generated UTRs demonstrated similar localized secondary structure profiles (Supplementary Fig. 3). The average resulting MFE traces are positively correlated between the experimental high-expression and designed high-expression UTRs, and negatively correlated between the designed high-performing and experimental low-performing sequences. This trend also holds when comparing the experimental low-expression and designed low-expression sequence groups, where a positive correlation is again observed, solidifying the ability of SANDSTORM-guided optimization to produce thermodynamically realistic sequences at both a local and global level. Despite being directly optimized with respect to function, the GARDN-designed UTRs can recreate the same distribution of secondary structure features as their experimental counterparts following activation maximization. These results suggest that during the predictive training process, meaningful abstractions of secondary structure are learned by the SANDSTORM model, and that these abstractions can effectively guide the exploration of the latent inputs to the generative model during the activation maximization progression.

### GARDN Modelling of Toehold Switches

Generative modelling of toehold switches is considerably more complex than 5’ UTRs due to the combination of specific sequence and structural motifs that define these devices. During preliminary tests, we found that unlike our GARDN-UTR generator, which leveraged self-attention layers, attention-based GARDN-Toehold generators struggled to adhere exactly to the target stem-loop structure required for toehold switch devices to function. Realistic toehold switch generation was accomplished by adapting a thematically similar approach to previous RNA inverse design software, where a target secondary structure can be specified by a user. This took the form of a reverse complementation layer in the generator network which operated on the latent nucleotide array, mapping the regions upstream and downstream of the RBS that base pair in the toehold switch before being passed through a final convolutional output layer. This development allowed several improvements offered through differentiable design while retaining the ability to design molecules with a target secondary structure.

We analyzed the resulting secondary structures of GARDN-generated toehold switches as well as those designed using activation maximization, constrained activation maximization (where the conserved sequence regions are included in the input seed)^47^, and NUPACK^13^. For a qualitative assessment of design quality, we computed the MFE structures of the toehold switch designs and overlaid them on top of one another to visualize the ensemble of secondary structures generated by the different design strategies (Fig. 5a-f). For quantitative assessment, rather than utilizing ensemble defect, which is only defined over the full sequence length, we employed a metric that could provide single-nucleotide resolution of structural agreement based on the expected paired or unpaired state of each nucleotide (see Methods). This metric was used to score how well the toehold switches designed using each algorithm adhered specifically to the canonical toehold switch structure (Fig. 5g,h). Sequences designed using NUPACK, without any added constraints on the target RNA sequence, showed the optimal structural distribution, as expected (Fig. 5a,h). The experimental toehold switch dataset used during the training process, which featured target RNA sequences from viruses and human genome sites, displayed lower structural agreement due to the additional sequence constraints but adopted the expected toehold switch secondary structure (Fig. 5b,h). In contrast, toehold switches designed using either standard or constrained activation maximization did not converge to a structure that would be expected to exhibit the key switching mechanism required to function (Fig. 5c,d), resulting in a significant reduction in structural agreement (Fig 5h). Designs generated using activation maximization also failed to recapitulate the conserved RBS and start codon sequences necessary for translation (Supplementary Fig. 4). GARDN-generated toehold switches, however, demonstrated a steady increase in structural agreement over adversarial training iterations, until finally converging to a value which replicates the distribution of secondary structures of the experimental toehold switch dataset used during the training process (Fig. 5g,h). We also used GARDN in tandem with the SANDSTORM-toehold predictor to generate toehold switches optimized to have extreme ON values. These GARDN-SANDSTORM designs not only retained the ensemble secondary structures of the training dataset during optimization, but additionally incorporated specific sequence preferences that matched those previously correlated with high ON-state expression in experiments based on thermodynamic considerations (Fig. 5f,h, Supplementary Fig. 5). These results demonstrate that a combination of the adversarial training setup and optimization guided by a structurally informed predictive model can allow for thermodynamically realistic, high performance sequences to be designed using GARDN models.

**Figure 5.**
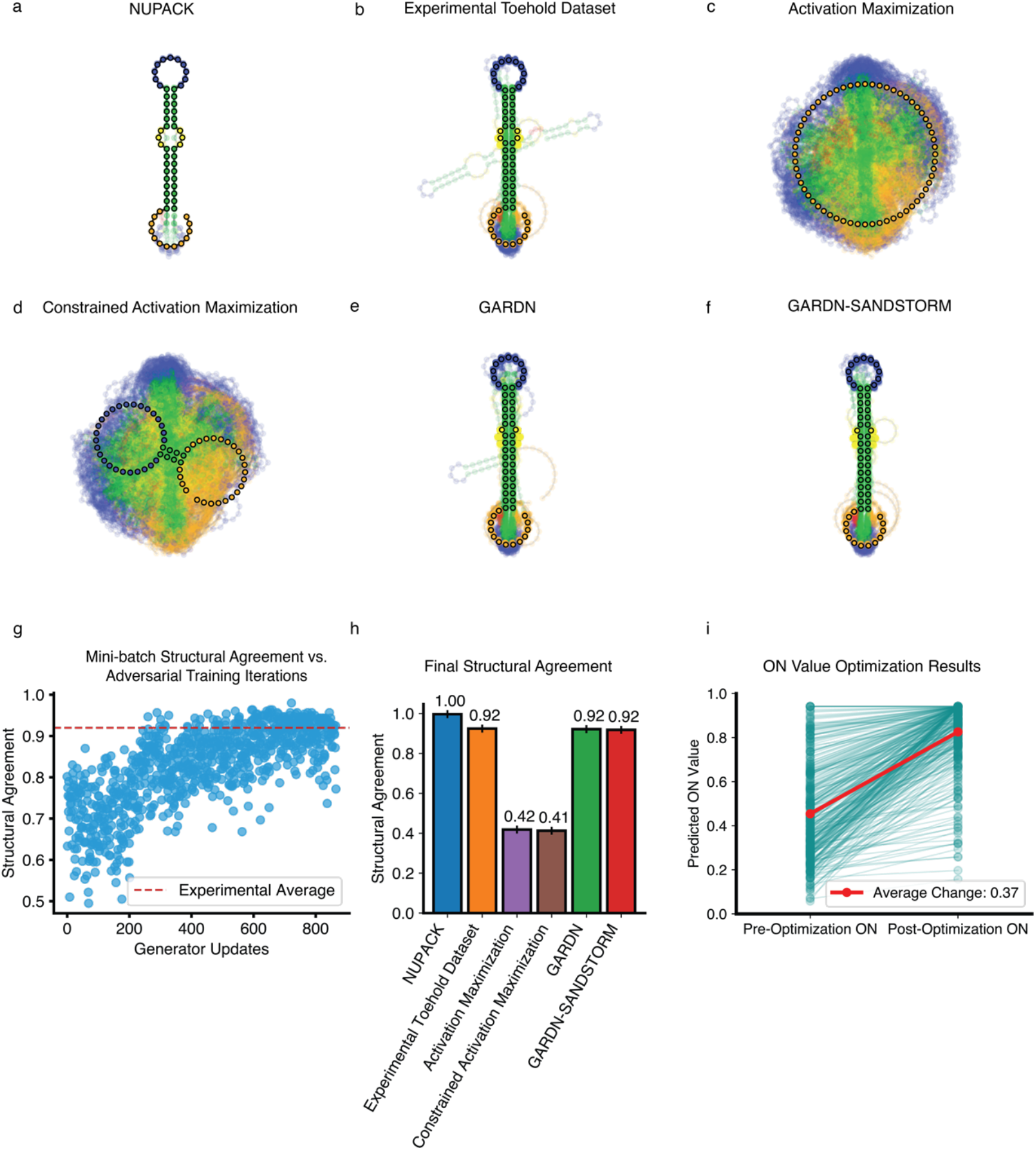
GARDN Models can design toehold switches with realistic secondary structures. **a-f**, The ensemble secondary structures of toehold switch sequences designed using a variety of algorithms, including those trained by experimental data: (a) NUPACK design algorithm applied without target RNA sequence constraints, (b) the high-throughput experimental dataset^33^ used for model training, (c) single sequence activation maximization, (d) single sequence activation maximization with starting seeds containing the constant motifs, (e) GARDN, and (f) GARDN-SANDSTORM with toehold switches optimized to have high ON values (n=300 a-f). Predicted structures were calculated using standard structural prediction software^13–15^, with the bolded structure representing the most likely structural character at each position (linearized for cases where the most likely structure is not valid). **g,** Structural agreement of GARDN-generated toehold switch sequences increases over adversarial training iterations, until converging to the experimental average. **h,** Final structural agreement between toehold switches designed using different algorithms and the canonical toehold switch secondary structure (quantification of a-f, see Methods). **i,** ON-value optimization results for GARDN-designed toehold switch sequences against the SANDSTORM-toehold predictive model. 300 calls to the predictive model resulted in a 37% increase in average ON score.

Previous approaches relying on post-hoc editing of designed sequences to achieve a plausible binding structure after scoring create a non-trivial disconnect between the model’s prediction of the sequence’s function before and after the adjustment. By optimizing a generator with an output that adheres to the expected grammar of a toehold switch, the latent space providing inputs to the GARDN models can be efficiently explored with respect to ON values without having to make manual edits to the resulting sequence that are destructive to the validity of the prediction (Fig. 5f). The structural agreement traces of both GARDN and GARDN-SANDSTORM constructs (Supplementary Fig. 5) reveal that these sequences deviate from the canonical toehold switch structure at the same loci as the experimental dataset used to train the generative model, a feature which is retained during functional optimization. While sequences designed using NUPACK showed the optimal structural distribution, this effect is expected as the GARDN models were trained to recreate the experimental distribution (Fig. 5b) encountered during training.

We found that a pre-trained GARDN model also demonstrates better nucleotide variability (Supplementary Fig. 4) and faster generation time (Supplementary Fig. 5) than standard inverse design algorithms^13–15^, with the added benefit of having a score predicting design function. This variability is critical in diagnostic and synthetic biology applications, where high overlap between multiple constructs can lead to unintended component cross-talk^3^. The lightweight SANDSTORM prediction architecture results in a simplified optimization landscape allowing for efficient sequence design. Using GARDN-SANDSTORM, we were able to achieve an average 37% percent increase in predicted normalized ON values of GARDN-generated sequences with only 300 calls to the predictive model (Fig. 5i). Improving the predicted scores of generated constructs was possible without sacrificing the realistic sequence contents of the resulting designs (Supplementary Fig. 4). Toehold switches optimized by GARDN-SANDSTORM demonstrated superior variation in the trigger and stem domains compared to other approaches while maintaining the constant RBS and start codon motifs. Moreover, comparing the sequence contents of GARDN-optimized toehold switches to those optimized with standard activation maximization and constrained activation maximization demonstrates that in the absence of a generator network to enforce realism, optimized sequences will trend towards solutions that trivially maximize the score of the predictor and sacrifice both sequence and structural realism (Fig. 5c,d).

### Experimental Validation of GARDN-SANDSTORM

To verify the performance improvement afforded by GARDN-SANDSTORM generative RNA design, we experimentally characterized the performance of several groups of toehold switch sequences designed using different algorithms. These groups consisted of non-optimized outputs from the GARDN models, GARDN sequences optimized to have high ON-state GFP expression using an ON-only SANDSTORM predictive model, GARDN sequences designed to have both a high ON state GFP expression and low OFF state GFP expression using a joint ON/OFF SANDSTORM model, and those designed using NUPACK (see Supplementary Tables 1 and 2 for sequences). The resulting designs were characterized in *E. coli* in conditions matching the toehold switch dataset used for training^33^ with the switch RNA on its own used to evaluate the OFF state and an RNA featuring a switch-trigger fusion used for the ON state (see Methods). Compared to the default GARDN output, sequences optimized with an ON-only SANDSTORM predictive model show an average 4.20-fold increase in GFP fluorescence from their parent sequences in the presence of the cognate trigger after only 300 calls to the predictive model (Fig. 6a). This result highlights the ability of our optimization process to improve specific attributes of RNA sequences.

**Figure 6.**
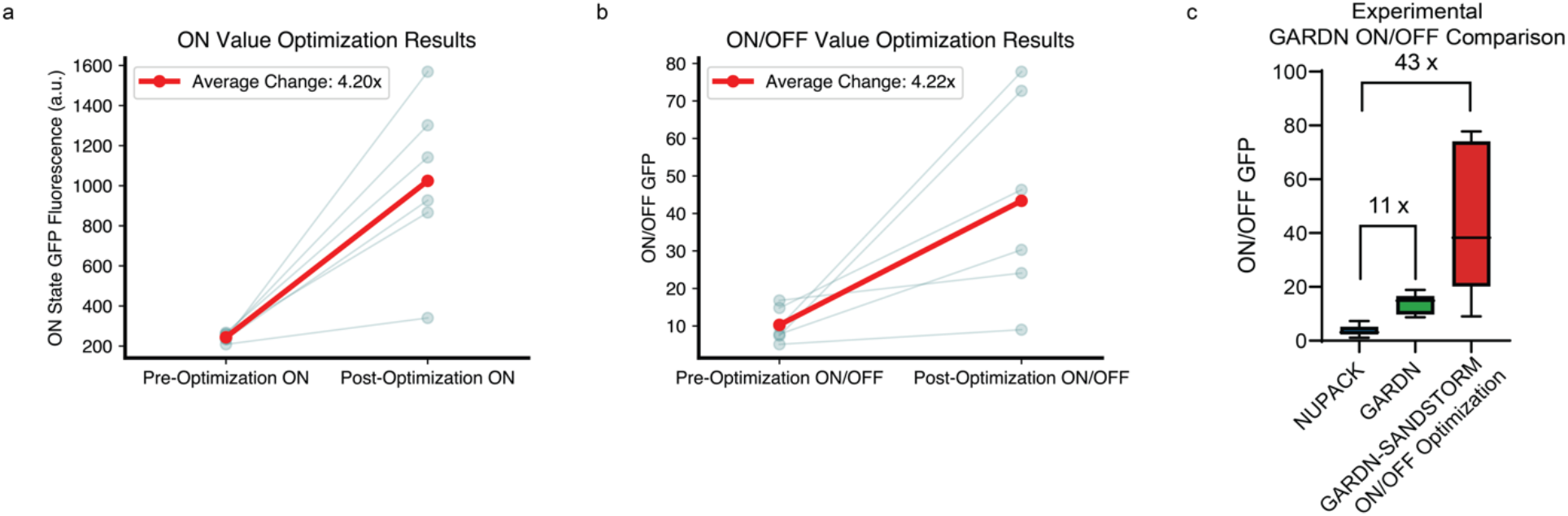
GARDN-designed toehold switches show improved experimental performance in *E. coli*. **a,** Toehold switch sequences designed by GARDN to have a high ON-state expression show a 4.20-fold increase in GFP expression in the presence of their cognate trigger in *E. coli* after optimization by a SANDSTORM predictive model (p = 0.0022, Mann-Whitney U test). **b,** SANDSTORM optimization of both ON- and OFF-state expression of GARDN sequences results in a 4.22-fold increase in ON/OFF ratio (p = 0.015, Mann-Whitney U test). **c,** Both non-optimized GARDN sequences and GARDN-SANDSTORM optimized groups show a significant increase in performance of 11-fold (p = 0.005, Mann-Whitney U test) and 43-fold (p = 0.005, Mann-Whitney U test), respectively, compared to those designed using classic inverse design software.

Optimization of ON/OFF ratios is a more complex task, as the predictor must convey meaningful information related to the performance of each sequence in two independent experimental states. Furthermore, the sequence or structural factors that minimize leakage in the OFF state, such as high GC content that creates a strong stem, are frequently at odds to those that maximize expression in the ON state, where an AU-rich stem can more readily unwind leading to increased expression^3^. Despite this dynamic tradeoff, GARDN-SANDSTORM optimization by a predictor trained on the ON/OFF ratios of toehold switch sequences demonstrated a 4.22-fold increase in ON/OFF ratios compared to their non-optimized counterparts (Fig. 6b). These findings demonstrate the capability of the GARDN-SANDSTORM approach to optimize RNA molecules with complex, multi-faceted functions, a key factor for the design of RNA across modern biotechnology settings.

Both the non-optimized GARDN and GARDN-SANDSTORM sequence groups demonstrated remarkable improvements of 11-fold and 43-fold, respectively, in experimental ON/OFF ratios compared to those designed using NUPACK (Fig. 6c). Importantly, this increase in performance does not come at the expense of either the sequence or structural realism of the designs.

## DISCUSSION

We have demonstrated that by incorporating the secondary structure of RNA molecules into a deep learning framework, we can achieve state-of-the-art functional prediction performance using a streamlined, generalizable convolutional neural network architecture. Additionally, we have shown that these models can extract interpretable abstractions of RNA structure – the presented generative models built around this encoding recreate experimentally validated thermodynamic properties while being optimized for function. These properties directly translate to improved experimental performance compared to current state-of-the-art sequence design tools while retaining critical conserved sequence elements, nucleotide diversity, and structural realism.

The GARDN approach to our knowledge offers the best available solution to designing structurally realistic RNA molecules with functions that cannot be accurately predicted using thermodynamic attributes alone. It is likely that this design paradigm shows improved performance by avoiding the underlying thermodynamic assumptions of standard inverse design tools while accounting for additional factors such as codon usage or mRNA-ribosome interactions that may be represented in high-throughput data. GARDN-SANDSTORM toehold switch optimization pairs additionally outperform the 2.6-fold improvement in ON/OFF ratios reported for activation maximization sequences from previous studies^33^, likely due to the elimination of manual sequence corrections.

Both SANDSTORM and GARDN models could be readily applied to new datasets mapping RNA variants to their functional performance. While our work focused on systems modulating translation efficiency and guide RNAs, other domains such as transcriptional riboregulators, IRES-like elements, aptamers, and RNA protein-binding recognition sites^54–58^ all have a critical dependency on RNA structure that would make them aptly suited for SANDSTORM predictions and GARDN design, given a suitable dataset. While there is no definitive lower bound on the number of sequences needed to deploy these models, the limited number of hyperparameters and generalized performance of SANDSTORM models suggest that fewer experimentally screened sequences could be used to train a SANDSTORM model versus other methods with a decreased risk of overfitting.

We expect that future work will focus on striking the same balance between biological-feature engineering and deep-learning-based feature abstraction that enable the enhanced performance of the SANDSTORM models. While the predictive capability demonstrated here could likely be improved by fine-tuning the dual-input method to each task, retaining generalizability is a critical constraint for the design of machine learning systems in biological contexts. The expansion of the latent reverse complementation layer of the GARDN-Toehold generator to an arbitrary target secondary structure could enable the benefits of differentiable design for several RNA devices, while the further development of the attention-based generators utilized for UTR design could allow the modelling of RNA molecules in settings with high structural heterogeneity.

As engineered functional RNAs continue to gain traction in various domains of biotechnology, the toolkit of computational methods for RNA design must keep pace. Reliable generative tools for engineering functional sequences will continue to be essential for rapidly adapting RNA systems to new use cases with minimal experimental screening. SANDSTORM and GARDN models offer a promising solution to the experimental screening challenge; the generalized ability to predict and design functional RNA molecules with increased performance while maintaining realistic secondary structures could serve as the foundation of a next-generation set of computational RNA tools.

## METHODS

### Secondary Structure Array Calculation

The secondary structure array developed in this work is an *N*×*N* array, where *N* is the length of an RNA sequence. Each point (*i*, *j*) in this array is assigned a value based on the possibility of the nucleotide at position *i* forming a Watson-Crick base pair with the nucleotide at position *j*. If (*i*, *j*) does not form a canonical base pair, a value of 0 is assigned. If (*i*, *j*) forms an A-U base pair, a value of 2 is assigned, and if (*i*, *j*) forms a G-C base pair, a value of 3 is assigned. These values reflect the number of hydrogen bonds that form in their respective base pairing interactions and are included to provide the model with information related to binding strength. Each position (*i*, *j*) is then divided by the distance along the sequence between nucleotide *i* and nucleotide *j* to encode positional information related to nucleotide interactions, and normalized row-wise to sum to 1.

### Structure-Based Classification Testing

Several classes of decoy toehold switches were constructed to evaluate the ability of the structural array to encode meaningful information. In this training setup, the dataset consisted of 8000 unique, one-hot-encoded sequences for each decoy class combined with the original toehold switch dataset from Angenent-Mari et al^33^. Standard multi-class classification was conducted on this synthetic dataset using a simplified version of both the two-input and sequence-only models and multi-categorical cross-entropy loss. The introduction of a second convolutional channel by default will increase the number of trainable parameters, which may lead to an artificial inflation in the performance of the two-channel models. To compensate for this, the two-input and sequence-only models were composed of the same depth of convolutional layers (3), but the number of filters per convolutional layer in the sequence-only model was increased to balance the number of trainable parameters between the two. The same split of data was used to compare the performance of the two models, with ROC curves and AUC metrics calculated using sklearn.

### SANDSTORM Predictive Architecture and Training

The full-sized predictive model consisted of two parallel stacks of three convolutional layers, with each convolutional stack extracting motifs from the one-hot-encoded sequence arrays and structural arrays, respectively. The outputs of these convolutions were concatenated before being passed through a series of three densely connected layers to a final output that returned a value for each functional property of the input sequences (e.g. ON and OFF for toehold switches, Mean Ribosome Loading for UTRs, etc.). Relu internal activation functions were used for all layers of the model and training was conducted using the Adam optimizer. The model was retrained for each specific dataset without hyperparameter tuning to ensure generalizability across classes. While hyperparameter tuning was not conducted, the resulting model size for each task does vary slightly due to the variation in size of the input sequence. Three-fold cross validation was used to benchmark against the previously published models, ensuring that the same data split was used for comparison at each iteration (see Supplementary Fig. 1 for a detailed diagram of the model architecture).

### GAN Framework

Generative adversarial networks consist of a generator model, *G*_θ_, and a discriminator model, *D*_φ_. The goal of the generator network is to produce novel examples of data that are indistinguishable from the real data distribution, *X*∼ℙ*_r_*, while the goal of the discriminator network is to differentiate between real and generated data samples. The generator network is a deep convolutional neural network parameterized by θ that accepts a continuous random variable input, *Z*∼ℙ*_z_*, sampled from a prior distribution. Data samples are generated by passing this random variable through several up sampling and convolutional layers to create an output, *G*_θ_(*Z*), that emulates the real data distribution. The discriminator model is also a convolutional neural network parameterized by φ that accepts both real and generated data and assigns a prediction to each that reflects which distribution they were sampled from, *D*_φ_(*G*_θ_(*Z*)) and *D*_φ_(*X*). These models simultaneously compete during the training process (standard gradient descent) until a generator network *G*_θ_ is realized that returns data samples that are indistinguishable from the original distribution.

A refined version of the GAN framework known as the Wasserstein GAN with gradient penalty (WGAN-GP)^51^ that is more robust to adjustments in hyperparameters was the foundation of the GARDN models. The key distinctions of this approach from standard GAN training involves altering the discriminator model to return a continuous realism score rather than a probability of the input coming from the real or generated class and the addition of a gradient penalty. This shifts the training process from minimizing the Jensen-Shannon divergence between the real and generated distributions to minimizing an estimation of the Earth Mover distance. In WGAN-GP models, weight clipping is replaced by a gradient penalty to enforce a soft version of the Lipschitz constraint. The gradient penalty is implemented by adding a term to the gradient norm of random uniform input samples 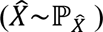 to the discriminator loss function:

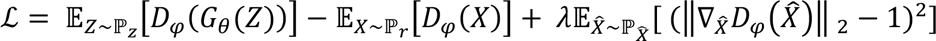

Where λ is the number of discriminator updates per generator update. To generate RNA sequences, we created GARDN models that accepted 128-dimensional gaussian vectors that were passed through several up-sampling and convolutional operations to return a 4×*N* array representing a single sequence of length *N*. The generative and discriminative components of the GARDN models were deep convolutional neural networks consisting of spectrally normalized convolutional layers with relu and leaky ReLU activation functions. For UTR generation, self-attention^53^ was additionally included in the internals of the models to allow for possible long-range nucleotide dependencies to take form. Hyperparameter tuning adjusting the size of convolutional filters, learning rates for each model, and number of discriminator updates per generator update was conducted until a stable training configuration that quickly converged to realistic sequences was achieved. Notably, the discriminator models used during GARDN training did not include the structural array input channel of the SANDSTORM models, as we found that doing so would result in a discriminator that converged too quickly relative to the generator for stable training to be achieved.

### GARDN Reverse Complementation Layer

A reverse complementation layer was developed to allow GARDN models to generate sequences adhering to the target structural grammar defining toehold switches. In this approach the one-hot-encoded values returned by the final up-sampling layer in the 5’ portion of the stem region of the toehold switch were reversed along their second (nucleotide) and third (sequence position) axes before being broadcast to the necessary 3’ portion of the stem region and passed through a final convolutional layer. This method greatly outperformed the addition of attention based internal layers at generating sequences with canonical toehold switch structures and in the future could be generalized to model arbitrary target RNA structures while maintaining valuable nucleotide diversity.

### GARDN Optimization

In a standard activation maximization setup, the input, *X*, to a differentiable predictive model, *P*_+_, is repeatedly summed with its own gradients to maximize the returned value while the parameters of the predictive model, δ, are held constant. This can be formalized as

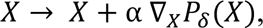

where α is a constant that is analogous to the learning rate of a standard neural network. In the case of sequence design, the inputs are a one-hot-encoded nucleotide array passed through some differentiable argmax operation (SoftMax, Gumbel SoftMax, Relaxed One-Hot Encoding) to preserve the discretized inputs expected by the predictive model while still allowing continuous gradient steps to take effect. To optimize GARDN outputs, we adapted the method from Killoran et al.^43^, where instead of directly optimizing the nucleotide array input to the predictive model, we take the gradient of the random variable seed, *Z*, of our generative model with respect to the predictive output, while freezing the weights of both the predictive and generative models (δ and θ). This approach allows us to move through the latent space providing input seeds to the generator until a region is found that returns sequences with desirable features as measured by the SANDSTORM predictor.

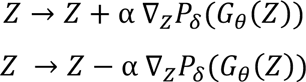

In this setup, the gradient step can either be summed with or subtracted from the original seed to move towards regions of the latent space that correspond to the generation of high- or low-performing sequences. At each gradient update step, the structural array input corresponding to the generated sequence is calculated and fed in parallel to the frozen SANDSTORM model; a key step that requires the memory and compute efficiencies of our novel structural array. For our study the predictor returned values related to the experimental function of RNA molecules; however, this predictor could be generalized to any arbitrary differentiable function.

### Computational Sequence Design Experiments

To design UTRs, we generated 300 high-expression UTR sequences using the gradient ascent setup (addition) and 300 low-expression sequences by using gradient descent (subtraction) on latent inputs to the generator. We compared the nucleotide contents and thermodynamic profiles of these generated constructs to the highest performing (mean ribosome loading > 8.5) and lowest performing (mean ribosome loading < 1.2) UTRs from the experimental dataset. For toehold switches, the predictive model returns two distinct outputs, one for ON and OFF. We chose to design 300 toehold switches with a maximized predicted ON value and compared to those designed by standard thermodynamic inverse design via NUPACK, single-sequence activation maximization, and single-sequence activation maximization with starting seeds that already contained the start codon and RBS motifs (constrained activation maximization). The GARDN-optimized sequences were designed using our novel two-input model, while both activation maximization approaches were designed by performing gradient ascent of a single sequence (n=300 total sequences) using the predictive model from Valeri et al^34^. The sequence logos demonstrated throughout Figs. 4 and 5 represent the average nucleotide frequencies of these groups of sequences.

### Experimental Library Design

A library of six toehold switch sequences were selected from several design approaches to validate the performance improvements offered through GARDN-SANDSTORM. To replicate the conditions of the high-throughput dataset used to train the SANDSTORM models, ON-state sequences were validated in the switch-trigger fusion configuration originally used by Angenet-Mari et. al^33^. For the most direct performance comparison to the previously published work, the SANDSTORM predictor shown in Figure 3 and the one used for the computational optimization in Figure 5 matched the formatting of the original work, where a single model predicts both the ON and OFF values of each sequence. In practice however, we found a substantial increase in performance by training on the predicted ON/OFF ratio itself as a single output, which we utilized to optimize the experimentally validated sequences. The first group consisted of six sequences generated randomly from GARDN outputs. Additionally, 100 GARDN sequences were optimized by an ON-only predictive SANDSTORM model (300 optimization calls), with the six sequences that demonstrated the highest predicted increase in ON state selected for experimental validation. This approach was repeated to select an additional six sequences with a SANDSTORM model that predicted the ON/OFF ratios. A final set of six sequences was designed using the inverse-design toolkit offered by NUPACK 4.0 under default model conditions^14^. For this group, inverse design was carried out by individually specifying the switch, trigger, and switch-trigger complexes without any sequence constraints beyond the RBS, start codon, and reporter gene sequences.

### Plasmid Construction

All sequences were ordered as single-stranded DNA oligonucleotides from Integrated DNA technologies with common 30 nucleotide homology regions. Following PCR to create double-stranded DNA, the amplified DNAs were then connected with plasmid backbones using Gibson assembly. Plasmid backbones were amplified from the commercially available pCOLADuet (EMD Millipore) expression vector via PCR followed by digestion with the restriction enzyme DpnI. All Gibson assembly products were transformed directly into BL21 DE3 cells, and sequence verified by Sanger sequencing. The reporter protein utilized in the study is GFPmut3b with an ASV degradation tag.

### Flow Cytometry Analysis

Bacterial colonies transformed with plasmids containing either the fused switch-trigger construct (ON state) or with just the unfused switch (OFF state) were inoculated in 1 mL of LB in triplicate with 50 µg mL^−1^ kanamycin antibiotic and grown overnight at 37°C shaking at 750 rpm. After 14 hours, 5 μL of overnight culture was diluted 100-fold in 495 μL of fresh LB with 25 µg mL^−1^ kanamycin. Following recovery until OD600 reached approximately 0.2, isopropyl β-D-1-thiogalactopyranoside (IPTG) was added to each well to a final concentration of 0.25 mM.

Following 3 hours of induction with IPTG, flow cytometry was performed with a Stratedigm S1000 cell analyzer equipped with an A600 Stratedigm high-throughput auto sampler. Prior to running flow cytometry, inoculated cultures were diluted 2-fold into phosphate buffered saline (PBS) in 384-well plates. Approximately 50,000 individual cells were recorded per sample, and cell populations were gated according to their forward side scatter (FSC) and side scatter (SSC) distributions to select for live cells and singlets, respectively, as described previously^3, 8^. The resulting mean GFP fluorescence signal generated by these gated populations was used for subsequent calculations. Fluorescence measurements of ON-state and OFF-state cells were calculated as the mean of three technical replicates, and error levels were calculated from the standard deviation of measurements from three biological replicates (nine samples total per construct).

### Thermodynamic Parameter Calculations

All results demonstrating the thermodynamic properties of sequences were calculated using NUPACK 4.0 in python under default model settings. We computed a sliding window minimum free energy calculation (Supplementary Fig. 3) by slicing out varying sizes of the sequences to plot traces of localized secondary structures and GC content. Distributions of ensemble defect are calculated across the entire sequence.

### Structural Agreement

In the case of toehold switches, a more specific quantification of secondary structure was needed to analyze the performance of the GARDN models than in the case of UTRs. To do so we employed the ‘structural agreement’ metric that reflects how well a dataset of generated sequences adheres to a specific secondary structure. This value is obtained by computing the predicted secondary structure of every sequence in a particular dataset in dot parens plus notation. In this notation, each position in a sequence can take the value of ‘.’ to represent an unpaired nucleotide, ‘(‘ to represent a nucleotide bound to a downstream nucleotide, and ‘)’ to represent a nucleotide bound to an upstream nucleotide. The frequencies of each of symbol at each position can be averaged across a dataset to form a consensus structural array representing the probability of finding each symbol at each position. The argmax can be computed along this array of structural probabilities to return the positional consensus secondary structure. For canonical toehold switches, the ideal secondary structure in dot parens plus notation is ‘…………(((((((((…((((((………..))))))…)))))))))’. We compute the structural agreement for a dataset of sequences by the following:

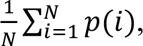

where *N* is the length of the canonical structure and *p*(*i*) is the probability of each canonical symbol appearing in the correct position as calculated by the consensus structural matrix. Without averaging, the values of *p*(*i*) can be plotted as a trace to provide single-nucleotide resolution of where a particular dataset diverges from the target structure (Supplementary Fig. 5).

## Acknowledgments

This work was supported by startup funds from Boston University to AAG. ATR was supported by the NIH Training Program in Quantitative Biology and Physiology (5T32GM008764). JMR was supported by the NIH Training Program in Synthetic Biology and Biotechnology (1T32GM130546) and a National Science Foundation Graduate Research Fellowship (2234657).

## Author contributions

ATR and AAG conceived of the overall idea and study design. ATR designed the models, implemented code, conducted data analysis, and wrote the manuscript. JMR conducted experimental validation of sensors and edited the manuscript. AAG edited the manuscript, supervised the project, and guided data analysis.

## Data Availability

The datasets used in this study can be found at the original source references.

