## Supplementary Information for "Generative and predictive neural networks for the design of functional RNA molecules"

### Supplementary Figures

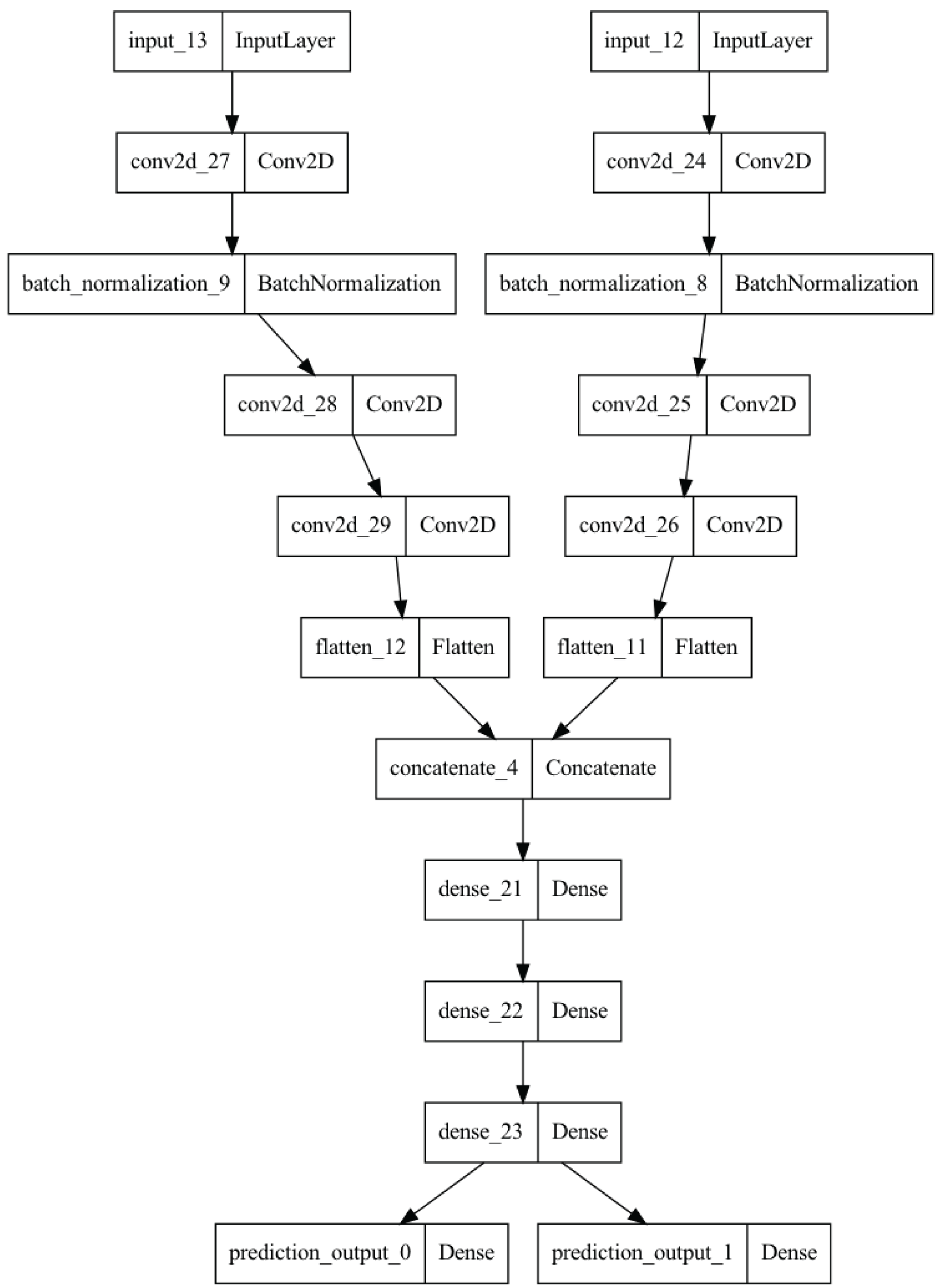


**Supplementary Figure 1 | SANDSTORM architecture.** Depicted is the architecture of the SANDSTORM models used to predict RNA function. The same model was used for the prediction of toehold switch ON/OFF values, UTR translation efficiency, RBS efficiency, and CRISPR gRNA/target efficiency, while re-training was conducted using the same hyperparameter combination on each dataset.

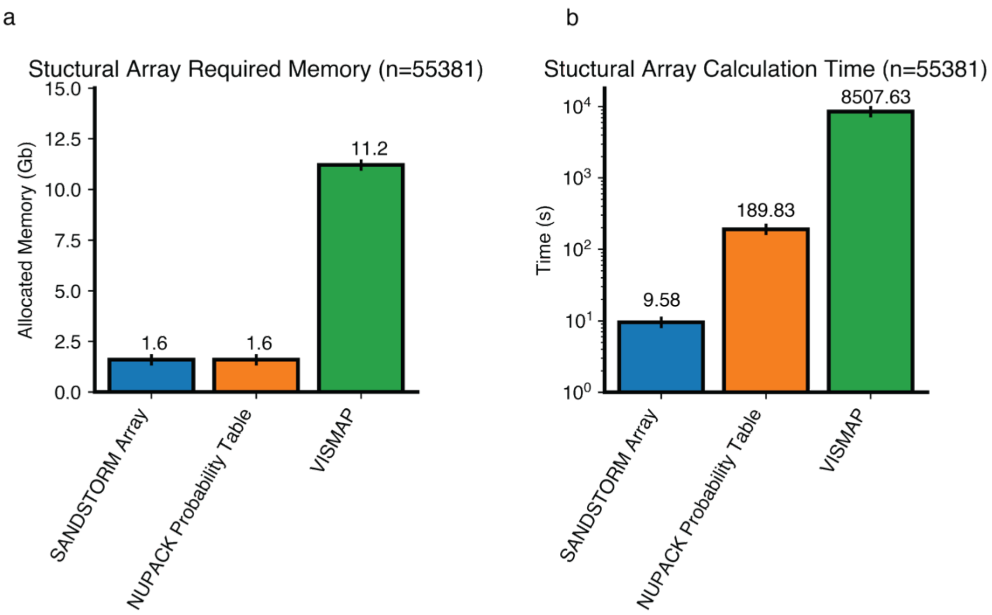


**Supplementary Figure 2 | Comparison of RNA structural array representations. a-b,** Two-dimensional representations of RNA secondary structure have been an essential component of computational RNA tools; however, the compute time, storage requirements, and thermodynamic assumptions of previously described array representations make them non-ideal for use in deep learning pipelines. Comparison of the required memory (a) and compute time (b) of the standard pairwise-probability array used by folding software based on thermodynamic models, such as NUPACK[1]; the Vismap representation of RNA structure published by Angenet-Mari et al.[2]; and the novel array used in this work.

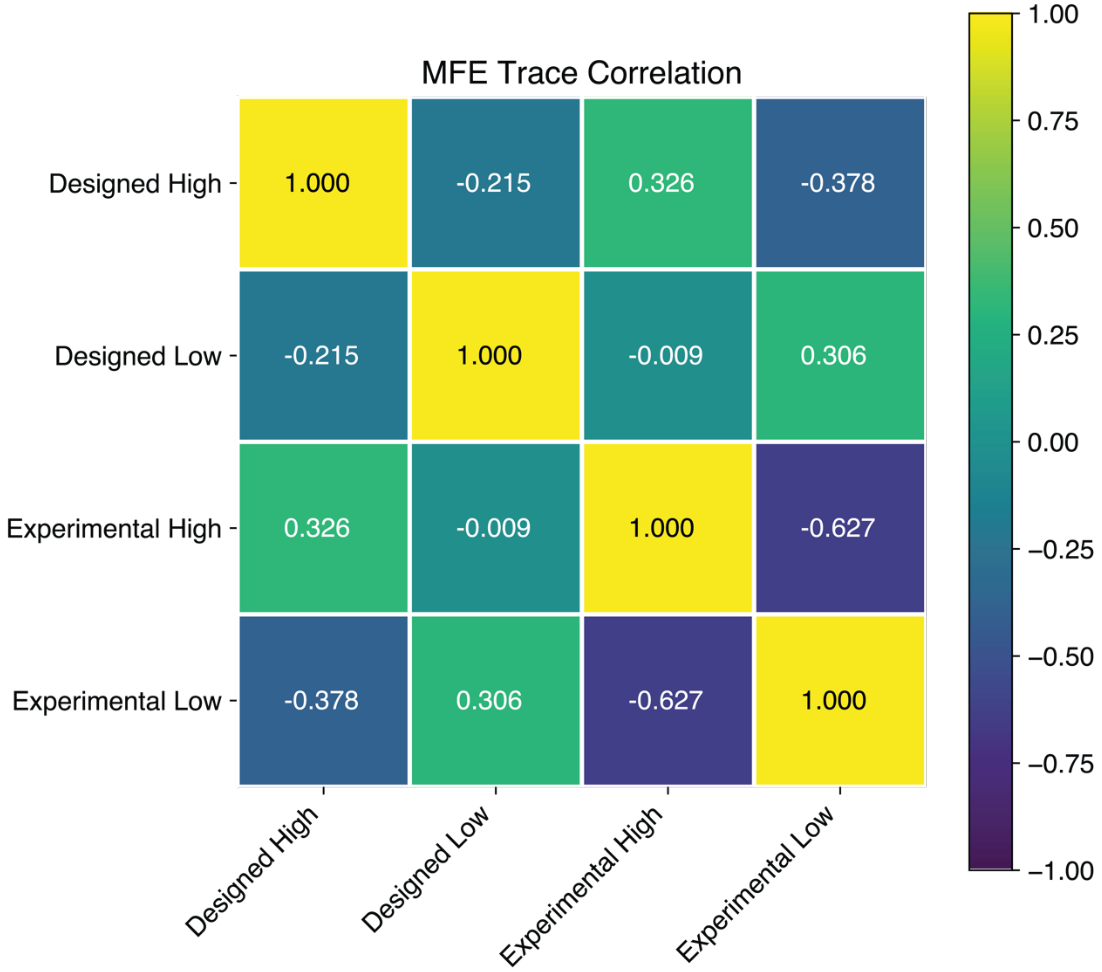


**Supplementary Figure 3 | Sliding window MFE correlations.** The localized secondary structure of UTR sequences was analyzed by selecting 20-nt sections of the sequence over a sliding window and using NUPACK[1] to calculate the MFE of each sequence section. This metric is of particular use for UTRs, where high increased structural complexity in proximity to the start codon could affect ribosomal progression and therefore resulting downstream translation efficiency[3-6].

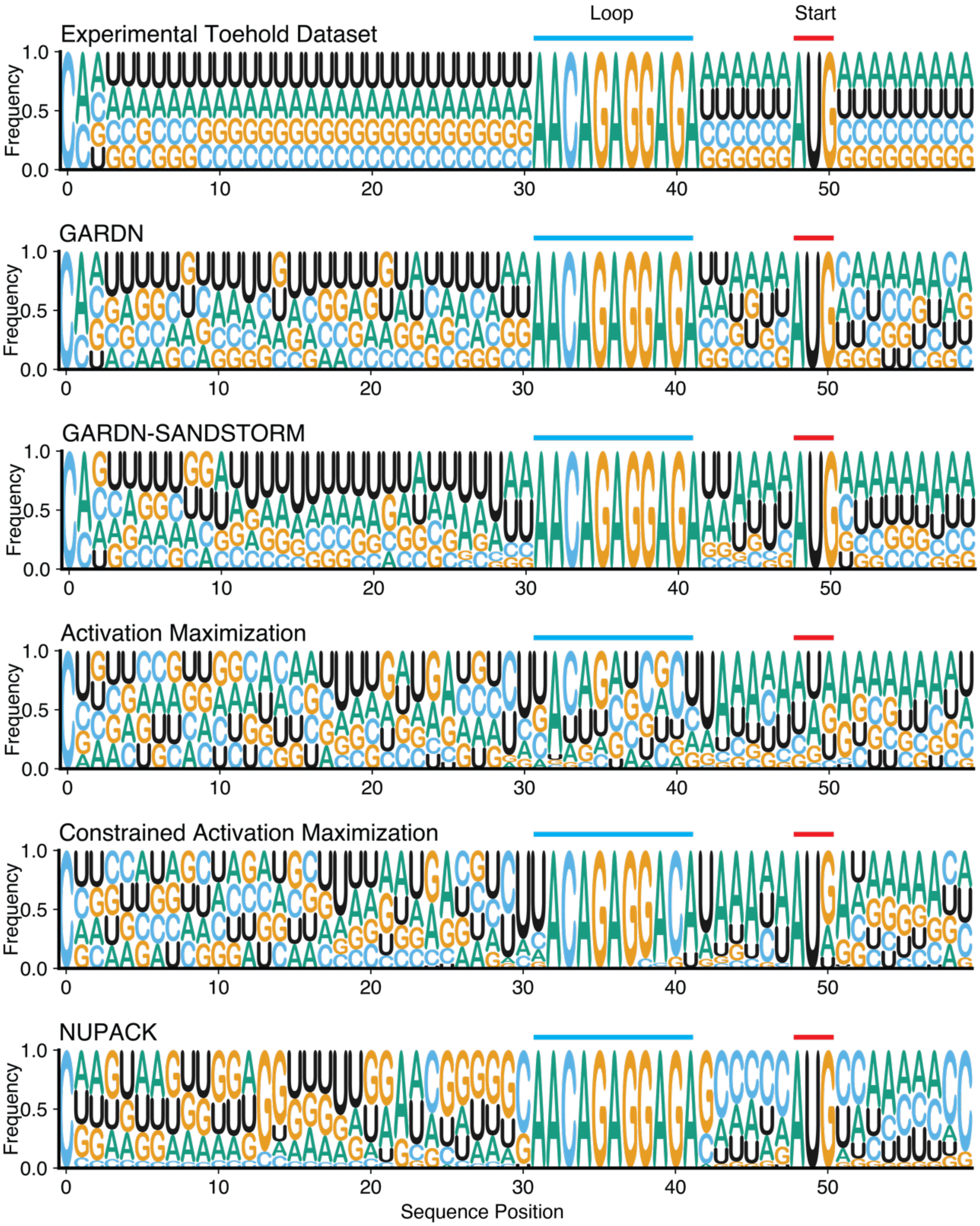


**Supplementary Figure 4 | Nucleotide distributions of toehold switch design algorithms.** The GARDN-generated and optimized toehold switches demonstrated superior nucleotide variability compared to the various RNA-design algorithms[1,7]. Activation maximization of a single sequence results in nucleotide patterns that maximize the predictive score while sacrificing variability. This effect can be seen specifically in the over-representation of A-U base pairs in the stem domain of activation maximization sequences, which would be expected to create a weak stem leading to high ON-state translation[8,9]. The GARDN-optimized sequences effectively incorporate this design rule, while also maintaining library diversity throughout the length of the entire construct. This effect is especially pronounced in the final two nucleotides of the stem domain, which has been previously correlated with high ON state expression[8]. NUPACK-based design of toehold switches results in strong G-C base pairing in the top of the stem, as these sequences are designed specifically against the secondary structure representation of a toehold switch, without incorporating function. The overrepresentation of G-C pairings in these sequences increase the strength of the stem and the likelihood of demonstrating the correct structure, while not necessarily improving function.

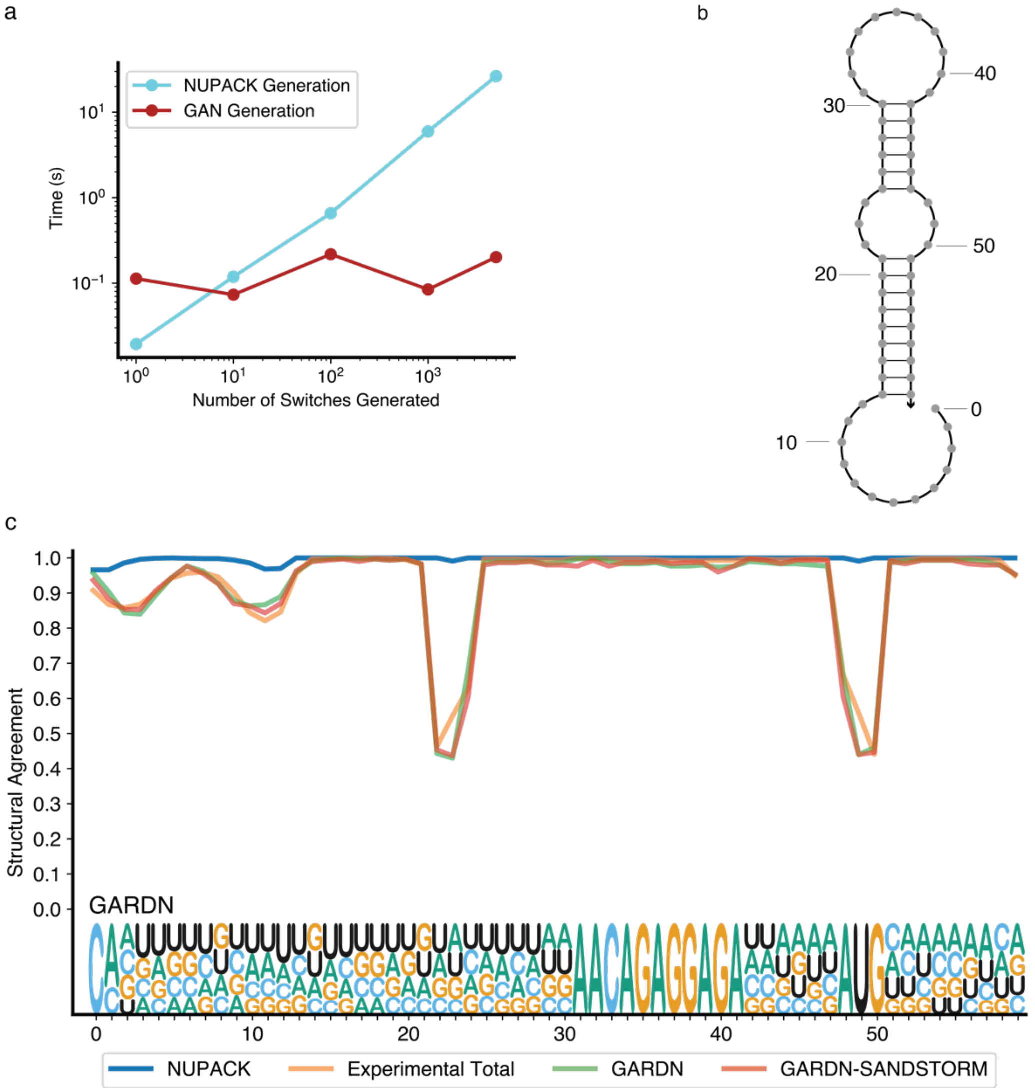


**Supplementary Figure 5 | Structural agreement over sequence length. a,** A pre-trained GARDN model demonstrates faster toehold-switch generation than standard inverse design algorithms (NUPACK[1]). **b,** Schematic of ideal toehold switch secondary structure along with numbered nucleotide positions. **c,** The secondary structure of toehold switch sequences designed using GARDN demonstrate variations from the canonical secondary structure at specific loci along the sequence that match the experimental training set, represented as dips in structural agreement. Notably, these adherent sections are not in regions affected by the reverse complementation layer enforcing the target structure but are learned throughout the adversarial training process. This trend holds in the GARDN-SANDSTORM sequence set as well, demonstrating that thermodynamic realism is enforced by the generative model when designing functional sequences.

### Supplementary Tables

**Supplementary Table 1 | Experimentally tested DNA constructs (switch-trigger fusions)**

| Name | Sequence |
|---|---|
| GARDN_0 | CTCTGGGCTAACTGTCGCGCTAATACGACTCACTATAGGGTTATTGTCAGTCCCTCAGCACACGCCGCCTAACCAAACACACAAACGCACAGGCGGCGTGTGCTGAGGGACTGACAATAAAACAGAGGAGATTATTGATGGTCCCTCAGAACCTGGCGGCAGCGCAAAAGATGCGTAAAGGAGAA |
| GARDN_1 | CTCTGGGCTAACTGTCGCGCTAATACGACTCACTATAGGGAATGAAGCCACTAAAGAAGCCACAAAAACTAACCAAACACACAAACGCACAGTTTTTGTGGCTTCTTTAGTGGCTTCATTAACAGAGGAGAAATGAAATGACTAAAGAAAACCTGGCGGCAGCGCAAAAGATGCGTAAAGGAGAA |
| GARDN_2 | CTCTGGGCTAACTGTCGCGCTAATACGACTCACTATAGGGGGTGAGCCTCCCGGTGAGGTCCTCGGGATTAACCAAACACACAAACGCACAATCCCGAGGACCTCACCGGGAGGCTCACCAACAGAGGAGAGGTGAGATGCCCGGTGAGAACCTGGCGGCAGCGCAAAAGATGCGTAAAGGAGAA |
| GARDN_3 | CTCTGGGCTAACTGTCGCGCTAATACGACTCACTATAGGGAAGACTAGAACTCCACTACATATCAATTATAACCAAACACACAAACGCACATAATTGATATGTAGTGGAGTTCTAGTCTTAACAGAGGAGAAAGACTATGACTCCACTAAACCTGGCGGCAGCGCAAAAGATGCGTAAAGGAGAA |
| GARDN_4 | CTCTGGGCTAACTGTCGCGCTAATACGACTCACTATAGGGTTCACAAATGTCGGAAGACATATAGGAGCTAACCAAACACACAAACGCACAGCTCCTATATGTCTTCCGACATTTGTGAAAACAGAGGAGATTCACAATGGTCGGAAGAAACCTGGCGGCAGCGCAAAAGATGCGTAAAGGAGAA |
| GARDN_5 | CTCTGGGCTAACTGTCGCGCTAATACGACTCACTATAGGGGGAATTAGCGACAACATTAGACAAAAGCAGAACCAAACACACAAACGCACCTGCTTTTGTCTAATGTTGTCGCTAATTCCAACAGAGGAGAGGAATTATGGACAACATTAACCTGGCGGCAGCGCAAAAGATGCGTAAAGGAGAA |
| ON_6_pre | CTCTGGGCTAACTGTCGCGCTAATACGACTCACTATAGGGCCATATGGGACGCCGGCCGACCACCCTGATAACCAAACACACAAACGCACATCAGGGTGGTCGGCCGGCGTCCCATATGGAACAGAGGAGACCATATATGACGCCGGCCAACCTGGCGGCAGCGCAAAAGATGCGTAAAGGAGAA |
| ON_6_post | CTCTGGGCTAACTGTCGCGCTAATACGACTCACTATAGGGAAATAATCAAAACCGAACATCCATTTCGATAACCAAACACACAAACGCACATCGAAATGGATGTTCGGTTTTGATTATTTAACAGAGGAGAAAATAAATGAAACCGAACAACCTGGCGGCAGCGCAAAAGATGCGTAAAGGAGAA |
| ON_7_pre | CTCTGGGCTAACTGTCGCGCTAATACGACTCACTATAGGGGGGCCGCCCCAAAGGTGGTGCAAGGCGATAACCAAACACACAAACGCACATCGCCTTGCACCACCTTTGGG |

| | |
|---|---|
| | GCGGCCCCAACAGAGGAGAGGGGCCATGCCAAAGGTGAACCTGGCGGCAGCGCA AAAGATGCGTAAAGGAGAA |
| ON_7_post | CTCTGGGCTAACTGTCGCGCTAATACGACTCACTATAGGGGGATTATCAACAAAGA TGATGCAAGGCCATAACCAAACACACAAACGCACATGGCCTTGCATCATCTTTGTT GATAATCCAACAGAGGAGAGGATTAATGACAAAGATGAACCTGGCGGCAGCGCAA AAGATGCGTAAAGGAGAA |
| ON_8_pre | CTCTGGGCTAACTGTCGCGCTAATACGACTCACTATAGGGCCGCCTAGAGCACCCC GGGTGTCATTTGTGAACCAAACACACAAACGCACCACAAATGACACCCGGGGTGC TCTAGGCGGAACAGAGGAGACCGCCTATGGCACCCCGGAACCTGGCGGCAGCGCA AAAGATGCGTAAAGGAGAA |
| ON_8_post | CTCTGGGCTAACTGTCGCGCTAATACGACTCACTATAGGGTTACCTCGACGACCTC GACTCTCATTAGTGAACCAAACACACAAACGCACCACTAATGAGAGTCGAGGTCG TCGAGGTAAAACAGAGGAGATTACCTATGCGACCTCGAAACCTGGCGGCAGCGCA AAAGATGCGTAAAGGAGAA |
| ON_9_pre | CTCTGGGCTAACTGTCGCGCTAATACGACTCACTATAGGGTTCTTCTTACGACCCAG GGTGCGGCCTCTTAACCAAACACACAAACGCACAAGAGGCCGCACCCTGGGTCGT AAGAAGAAAACAGAGGAGATTCTTCATGCGACCCAGGAACCTGGCGGCAGCGCAA AAGATGCGTAAAGGAGAA |
| ON_9_post | CTCTGGGCTAACTGTCGCGCTAATACGACTCACTATAGGGTTATTATTAGGACCAC TGCTGCTATTCCTTAACCAAACACACAAACGCACAAGGAATAGCAGCAGTGGTCCT AATAATAAAACAGAGGAGATTATTAATGGGACCACTGAACCTGGCGGCAGCGCAA AAGATGCGTAAAGGAGAA |
| ON_10_pre | CTCTGGGCTAACTGTCGCGCTAATACGACTCACTATAGGGCCGAACACTTCCCCAA AGGTGTGAAATTTTAACCAAACACACAAACGCACAAAATTTCACACCTTTGGGGAA GTGTTCGGAACAGAGGAGACCGAACATGTCCCCAAAGAACCTGGCGGCAGCGCAA AAGATGCGTAAAGGAGAA |
| ON_10_post | CTCTGGGCTAACTGTCGCGCTAATACGACTCACTATAGGGTTAAAATCTTCTCCAA ACATTCCAGGCTTTAACCAAACACACAAACGCACAAAGCCTGGAATGTTTGGAGA AGATTTTAAAACAGAGGAGATTAAAAATGTCTCCAAACAACCTGGCGGCAGCGCA AAAGATGCGTAAAGGAGAA |
| ON_11_pre | CTCTGGGCTAACTGTCGCGCTAATACGACTCACTATAGGGGGTCGCAACGGAGGAT ATTCAACAGGGCCTAACCAAACACACAAACGCACAGGCCCTGTTGAATATCCTCCG TTGCGACCAACAGAGGAGAGGTCGCATGGGAGGATATAACCTGGCGGCAGCGCAA AAGATGCGTAAAGGAGAA |
| ON_11_post | CTCTGGGCTAACTGTCGCGCTAATACGACTCACTATAGGGGGACTACACAGAGGAT ACTCAGCAGGCCCTAACCAAACACACAAACGCACAGGGCCTGCTGAGTATCCTCTG TGTAGTCCAACAGAGGAGAGGACTAATGAGAGGATACAACCTGGCGGCAGCGCAA AAGATGCGTAAAGGAGAA |
| ON_OFF_12_ pre | CTCTGGGCTAACTGTCGCGCTAATACGACTCACTATAGGGCCTGCTAGACCTCCGG GAGATCCTTTCGCTAACCAAACACACAAACGCACAGCGAAAGGATCTCCCGGAGG TCTAGCAGGAACAGAGGAGACCTGCTATGCCTCCGGGAAACCTGGCGGCAGCGCA AAAGATGCGTAAAGGAGAA |
| ON_OFF_12_ post | CTCTGGGCTAACTGTCGCGCTAATACGACTCACTATAGGGTTGGTTTGACCTAAAA GAAATTTGAACTCTAACCAAACACACAAACGCACAGAGTTCAAATTTCTTTTAGGT CAAACCAAAACAGAGGAGATTGGTTATGCCTAAAAGAAACCTGGCGGCAGCGCAA AAGATGCGTAAAGGAGAA |
| ON_OFF_13_ pre | CTCTGGGCTAACTGTCGCGCTAATACGACTCACTATAGGGCCGCATCGTATCCCAC AGCACATAAAGTCTAACCAAACACACAAACGCACAGACTTTATGTGCTGTGGGAT |

| | |
|---|---|
| | ACGATGCGGAACAGAGGAGACCGCATATGATCCCACAGAACCTGGCGGCAGCGCAAAAGATGCGTAAAGGAGAA |
| ON_OFF_13_post | CTCTGGGCTAACTGTCGCGCTAATACGACTCACTATAGGGCCATATTGAATCAAACGGCACATAAAGTGGAACCAAACACACAAACGCACCCACTTTATGTGCCGTTTGATTCAATATGGAACAGAGGAGACCATATATGATCAAACGGAACCTGGCGGCAGCGCAAAAGATGCGTAAAGGAGAA |
| ON_OFF_14_pre | CTCTGGGCTAACTGTCGCGCTAATACGACTCACTATAGGGCCAGAGGACTGCAACAGGATAAAAGGTCTTAACCAAACACACAAACGCACAAGACCTTTTATCCTGTTGCAGTCCTCTGGAACAGAGGAGACCAGAGATGTGCAACAGGAACCTGGCGGCAGCGCAAAAGATGCGTAAAGGAGAA |
| ON_OFF_14_post | CTCTGGGCTAACTGTCGCGCTAATACGACTCACTATAGGGTTAGAGGATTGCAACAGGATGTAGGGTCTTAACCAAACACACAAACGCACAAGACCCTACATCCTGTTGCAATCCTCTAAAACAGAGGAGATTAGAGATGTGCAACAGGAACCTGGCGGCAGCGCAAAAGATGCGTAAAGGAGAA |
| ON_OFF_15_pre | CTCTGGGCTAACTGTCGCGCTAATACGACTCACTATAGGGCCCAAGATGATTTTAACAAATATTAAGCTTAACCAAACACACAAACGCACAAGCTTAATATTTGTTAAAATCATCTTGGGAACAGAGGAGACCCAAGATGATTTTAACAAACCTGGCGGCAGCGCAAAAGATGCGTAAAGGAGAA |
| ON_OFF_15_post | CTCTGGGCTAACTGTCGCGCTAATACGACTCACTATAGGGAACAAGATGATTGGAGCGGCGATAAAGCTTAACCAAACACACAAACGCACAAGCTTTATCGCCGCTCCAATCATCTTGTTAACAGAGGAGAAACAAGATGATTGGAGCGAACCTGGCGGCAGCGCAAAAGATGCGTAAAGGAGAA |
| ON_OFF_16_pre | CTCTGGGCTAACTGTCGCGCTAATACGACTCACTATAGGGTTCACTAATGGGGGCCAGCATCGGGGAACTAACCAAACACACAAACGCACAGTTCCCCGATGCTGGCCCCCATTAGTGAAAACAGAGGAGATTCACTATGGGGGGCCAGAACCTGGCGGCAGCGCAAAAGATGCGTAAAGGAGAA |
| ON_OFF_16_post | CTCTGGGCTAACTGTCGCGCTAATACGACTCACTATAGGGTTCAGTAATAGAGGGGTAGATCTGGGAGCTAACCAAACACACAAACGCACAGCTCCCAGATCTACCCCTCTATTACTGAAAACAGAGGAGATTCAGTATGAGAGGGGTAAACCTGGCGGCAGCGCAAAAGATGCGTAAAGGAGAA |
| ON_OFF_17_pre | CTCTGGGCTAACTGTCGCGCTAATACGACTCACTATAGGGCCATTGTACGGGCCGAACAACCCCCCCAGTAACCAAACACACAAACGCACACTGGGGGGGTTGTTCGGCCCGTACAATGGAACAGAGGAGACCATTGATGGGGCCGAACAACCTGGCGGCAGCGCAAAAGATGCGTAAAGGAGAA |
| ON_OFF_17_post | CTCTGGGCTAACTGTCGCGCTAATACGACTCACTATAGGGTTATTTAAGGGGGGACTAGACCTCCATGGAACCAAACACACAAACGCACCCATGGAGGTCTAGTCCCCCCTTAAAATAAAACAGAGGAGATTATTTATGGGGGGACTAACCTGGCGGCAGCGCAAAAGATGCGTAAAGGAGAA |
| NUPACK_0 | CTCTGGGCTAACTGTCGCGCTAATACGACTCACTATAGGGGCCTTCGACTCACTTACCTTTTTATTTTTCAACCAAACACACAAACGCACGAAAAATAAAAAGGTAAGTGAGTCGAAGGCAACAGAGGAGAGCCTTCATGTCACTTACCAACCTGGCGGCAGCGCAAAAGATGCGTAAAGGAGAA |
| NUPACK_1 | CTCTGGGCTAACTGTCGCGCTAATACGACTCACTATAGGGCCACCGCTTCATACTACCTCCCCCTACTACAACCAAACACACAAACGCACGTAGTAGGGGGAGGTAGTATGAAGCGGTGGAACAGAGGAGACCACCGATGCATACTACCAACCTGGCGGCAGCGCAAAAGATGCGTAAAGGAGAA |
| NUPACK_2 | CTCTGGGCTAACTGTCGCGCTAATACGACTCACTATAGGGCATCCAGACTCAAAAACCCACTACCCCCTTAACCAAACACACAAACGCACAAGGGGGTAGTGGGTTTTTGAG |

| | |
|---|---|
| | TCTGGATGAACAGAGGAGACATCCAATGTCAAAAACCAACCTGGCGGCAGCGCAA AAGATGCGTAAAGGAGAA |
| NUPACK_3 | CTCTGGGCTAACTGTCGCGCTAATACGACTCACTATAGGGGGTAAGGGTGAAAGTG AGTAGGAGAAGGGGAACCAAACACACAAACGCACCCCCTTCTCCTACTCACTTTCA CCCTTACCAACAGAGGAGAGGTAAGATGGAAAGTGAGAACCTGGCGGCAGCGCAA AAGATGCGTAAAGGAGAA |
| NUPACK_4 | CTCTGGGCTAACTGTCGCGCTAATACGACTCACTATAGGGGTCCAGCCTCTTTCCCA CCCATCCATATCTAACCAAACACACAAACGCACAGATATGGATGGGTGGGAAAGA GGCTGGACAACAGAGGAGAGTCCAGATGCTTTCCCACAACCTGGCGGCAGCGCAA AAGATGCGTAAAGGAGAA |
| NUPACK_5 | CTCTGGGCTAACTGTCGCGCTAATACGACTCACTATAGGGCGCCACCTTCCAATAC CCACAACCCTAACCAACCAAACACACAAACGCACGGTTAGGGTTGTGGGTATTGG AAGGTGGCGAACAGAGGAGACGCCACATGCCAATACCCAACCTGGCGGCAGCGCA AAAGATGCGTAAAGGAGAA |

### Supplementary Table 2 | Experimentally verified RNA Sequences

| Name | Trigger | Switch |
|---|---|---|
| GARDN_0 | UUAUUGUCAGUCCCU CAGCACACGCCGCCU | CAGGCGGCGUGUGCUGAGGGACUGACAAUAAAACAGA GGAGAUUAUUGAUGGUCCCUCAG |
| GARDN_1 | AAUGAAGCCACUAAA GAAGCCACAAAAACU | CAGUUUUUGUGGCUUCUUUAGUGGCUUCAUUAACAGA GGAGAAAUGAAAUGACUAAAGAA |
| GARDN_2 | GGUGAGCCUCCCGGU GAGGUCCUCGGGAUU | CAAUCCCGAGGACCUCACCGGGAGGCUCACCAACAGAG GAGAGGUGAGAUGCCCGGUGAG |
| GARDN_3 | AAGACUAGAACUCCA CUACAUAUCAAUUAU | CAUAAUUGAUAUGUAGUGGAGUUCUAGUCUUAACAGA GGAGAAAGACUAUGACUCCACUA |
| GARDN_4 | UUCACAAAUGUCGGA AGACAUAUAGGAGCU | CAGCUCCUAUAUGUCUUCCGACAUUUGUGAAAACAGA GGAGAUUCACAAUGGUCGGAAGA |
| GARDN_5 | GGAAUUAGCGACAAC AUUAGACAAAAGCAG | CCUGCUUUUGUCUAAUGUUGUCGCUAAUUCCAACAGA GGAGAGGAAUUAUGGACAACAUU |
| ON_6_pre | CCAUAUGGGACGCCG GCCGACCACCCUGAU | CAUCAGGGUGGUCGGCCGGCGUCCCAUAUGGAACAGA GGAGACCAUAUAUGACGCCGGCC |
| ON_6_post | AAAUAAUCAAAACCG AACAUCCAUUUCGAU | CAUCGAAAUGGAUGUUCGGUUUUGAUUAUUUAACAGA GGAGAAAAUAAAUGAAACCGAAC |
| ON_7_pre | GGGGCCGCCCCAAAG GUGGUGCAAGGCGAU | CAUCGCCUUGCACCACCUUUGGGGCGGCCCCAACAGAG GAGAGGGGCCAUGCCAAAGGUG |
| ON_7_post | GGAUUAUCAACAAAG AUGAUGCAAGGCCAU | CAUGGCCUUGCAUCAUCUUUGUUGAUAAUCCAACAGA GGAGAGGAUUAAUGACAAAGAUG |
| ON_8_pre | CCGCCUAGAGCACCCC GGGUGUCAUUUGUG | CCACAAAUGACACCCGGGGUGCUCUAGGCGGAACAGA GGAGACCGCCUAUGGCACCCCGG |
| ON_8_post | UUACCUCGACGACCU CGACUCUCAUUAGUG | CCACUAAUGAGAGUCGAGGUCGUCGAGGUAAAACAGA GGAGAUUACCUAUGCGACCUCGA |
| ON_9_pre | UUCUUCUUACGACCC AGGGUGCGGCCUCUU | CAAGAGGCCGCACCCUGGGUCGUAAGAAGAAAACAGA GGAGAUUCUUCAUGCGACCCAGG |
| ON_9_post | UUAUUAUUAGGACCA CUGCUGCUAUUCCUU | CAAGGAAUAGCAGCAGUGGUCCUAAUAAUAAAACAGA GGAGAUUAUUAAUGGGACCACUG |
| ON_10_pre | CCGAACACUUCCCCA AAGGUGUGAAAUUUU | CAAAAUUUCACACCUUUGGGGAAGUGUUCGGAACAGA GGAGACCGAACAUGUCCCCAAAG |

| | | |
|---|---|---|
| ON_10_post | UUAAAAUCUUCUCCAAACAUUCCAGGCUUU | CAAAGCCUGGAAUGUUUGGAGAAGAUUUUAAAACAGAGGAGAUUAAAAAUGUCUCCAAAC |
| ON_11_pre | GGUCGCAACGGAGGAUAUUCAACAGGGCCU | CAGGCCCUGUUGAAUAUCCUCCGUUGCGACCAACAGAGGAGAGGUCGCAUGGGAGGAUAU |
| ON_11_post | GGACUACACAGAGGAUACUCAGCAGGCCCU | CAGGGCCUGCUGAGUAUCCUCUGUGUAGUCCAACAGAGGAGAGGACUAAUGAGAGGAUAC |
| ON_OFF_12_pre | CCUGCUAGACCUCCGGGAGAUCCUUUCGCU | CAGCGAAAGGAUCUCCCGGAGGUCUAGCAGGAACAGAGGAGACCUGCUAUGCCUCCGGGA |
| ON_OFF_12_post | UUGGUUUGACCUAAAAGAAAUUUGAACUCU | CAGAGUUCAAAUUUCUUUUAGGUCAAACCAAAACAGAGGAGAUUGGUUAUGCCUAAAAGA |
| ON_OFF_13_pre | CCGCAUCGUAUCCCACAGCACAUAAAGUCU | CAGACUUUAUGUGCUGUGGGAUACGAUGCGGAACAGAGGAGACCGCAUAUGAUCCCACAG |
| ON_OFF_13_post | CCAUAUUGAAUCAAACGGCACAUAAAGUGG | CCCACUUUAUGUGCCGUUUGAUUCAAUAUGGAACAGAGGAGACCAUAUAUGAUCAAACGG |
| ON_OFF_14_pre | CCAGAGGACUGCAACAGGAUAAAAGGUCUU | CAAGACCUUUUAUCCUGUUGCAGUCCUCUGGAACAGAGGAGACCAGAGAUGUGCAACAGG |
| ON_OFF_14_post | UUAGAGGAUUGCAACAGGAUGUAGGGUCUU | CAAGACCCUACAUCCUGUUGCAAUCCUCUAAAACAGAGGAGAUUAGAGAUGUGCAACAGG |
| ON_OFF_15_pre | CCCAAGAUGAUUUUAACAAAUAUUAAGCUU | CAAGCUUAAUAUUUGUUAAAAUCAUCUUGGGAACAGAGGAGACCCAAGAUGAUUUUAACA |
| ON_OFF_15_post | AACAAGAUGAUUGGAGCGGCGAUAAAGCUU | CAAGCUUUAUCGCCGCUCCAAUCAUCUUGUUAACAGAGGAGAAACAAGAUGAUUGGAGCG |
| ON_OFF_16_pre | UUCACUAAUGGGGGCCAGCAUCGGGGAACU | CAGUUCCCCGAUGCUGGCCCCCAUUAGUGAAAACAGAGGAGAUUCACUAUGGGGGGCCAG |
| ON_OFF_16_post | UUCAGUAAUAGAGGGGUAGAUCUGGGAGCU | CAGCUCCCAGAUCUACCCCUCUAUUACUGAAAACAGAGGAGAUUCAGUAUGAGAGGGGUA |
| ON_OFF_17_pre | CCAUUGUACGGGCCGAACAACCCCCCCAGU | CACUGGGGGGGUUGUUCGGCCCGUACAAUGGAACAGAGGAGACCAUUGAUGGGGCCGAAC |
| ON_OFF_17_post | UUAUUUUAAGGGGGGACUAGACCUCCAUGG | CCCAUGGAGGUCUAGUCCCCCCUUAAAAUAAAACAGAGGAGAUUAUUUAUGGGGGGGACU |
| NUPACK_0 | GCCUUCGACUCACUUACCUUUUUAUUUUUC | CGAAAAAUAAAAAGGUAAGUGAGUCGAAGGCAACAGAGGAGAGCCUUCAUGUCACUUACC |
| NUPACK_1 | CCACCGCUUCAUACUACCUCCCCCUACUAC | CGUAGUAGGGGGAGGUAGUAUGAAGCGGUGGAACAGAGGAGACCACCGAUGCAUACUACC |
| NUPACK_2 | CAUCCAGACUCAAAAACCCACUACCCCCUU | CAAGGGGGUAGUGGGUUUUUGAGUCUGGAUGAACAGAGGAGACAUCCAAUGUCAAAAACC |
| NUPACK_3 | GGUAAGGGUGAAAGUGAGUAGGAGAAGGGG | CCCCCUUCUCCUACUCACUUUCACCCUUACCAACAGAGGAGAGGUAAGAUGGAAAGUGAG |
| NUPACK_4 | GUCCAGCCUCUUUCCCACCCAUCCAUAUCU | CAGAUAUGGAUGGGUGGGAAAGAGGCUGGACAACAGAGGAGAGUCCAGAUGCUUUCCCAC |
| NUPACK_5 | CGCCACCUUCCAAUACCCACAACCCUAACC | CGGUUAGGGUUGUGGGUAUUGGAAGGUGGCGAACAGAGGAGACGCCACAUGCCAAUACCC |
